# Mutation-specific pathophysiological mechanisms define different neurodevelopmental disorders associated with SATB1 dysfunction

**DOI:** 10.1101/2020.10.23.352278

**Authors:** J. den Hoed, E. de Boer, N. Voisin, A.J.M. Dingemans, N. Guex, L. Wiel, C. Nellaker, S.M. Amudhavalli, S. Banka, F.S. Bena, B. Ben-Zeev, V.R. Bonagura, A.-L. Bruel, T. Brunet, H.G. Brunner, H.B. Chew, J. Chrast, L. Cimbalistienė, H. Coon, The DDD study, E.C. Délot, F. Démurger, A.-S. Denommé-Pichon, C. Depienne, D. Donnai, D.A. Dyment, O. Elpeleg, L. Faivre, C. Gilissen, L. Granger, B. Haber, Y. Hachiya, Y. Hamzavi Abedi, J. Hanebeck, J.Y. Hehir-Kwa, B. Horist, T. Itai, A. Jackson, R. Jewell, K.L. Jones, S. Joss, H. Kashii, M. Kato, A.A. Kattentidt-Mouravieva, F. Kok, U. Kotzaeridou, V. Krishnamurthy, V. Kučinskas, A. Kuechler, A. Lavillaureix, P. Liu, L. Manwaring, N. Matsumoto, B. Mazel, K. McWalter, V. Meiner, M.A. Mikati, S. Miyatake, T. Mizuguchi, L.H. Moey, S. Mohammed, H. Mor-Shaked, H. Mountford, R. Newbury-Ecob, S. Odent, L. Orec, M. Osmond, T.B. Palculict, M. Parker, A. Petersen, R. Pfundt, E. Preikšaitienė, K. Radtke, E. Ranza, J.A. Rosenfeld, T. Santiago-Sim, C. Schwager, M. Sinnema, L. Snijders Blok, R.C. Spillmann, A.P.A. Stegmann, I. Thiffault, L. Tran, A. Vaknin-Dembinsky, J.H. Vedovato-dos-Santos, S.A. Vergano, E. Vilain, A. Vitobello, M. Wagner, A. Waheeb, M. Willing, B. Zuccarelli, U. Kini, D.F. Newbury, T. Kleefstra, A. Reymond, S.E. Fisher, L.E.L.M. Vissers

## Abstract

Whereas large-scale statistical analyses can robustly identify disease-gene relationships, they do not accurately capture genotype-phenotype correlations or disease mechanisms. We use multiple lines of independent evidence to show that different variant types in a single gene, *SATB1*, cause clinically overlapping but distinct neurodevelopmental disorders. Clinical evaluation of 42 individuals carrying *SATB1* variants identified overt genotype-phenotype relationships, associated with different pathophysiological mechanisms, established by functional assays. Missense variants in the CUT1 and CUT2 DNA-binding domains result in stronger chromatin binding, increased transcriptional repression and a severe phenotype. Contrastingly, variants predicted to result in haploinsufficiency are associated with a milder clinical presentation. A similarly mild phenotype is observed for individuals with premature protein truncating variants that escape nonsense-mediated decay and encode truncated proteins, which are transcriptionally active but mislocalized in the cell. Our results suggest that in-depth mutation-specific genotype-phenotype studies are essential to capture full disease complexity and to explain phenotypic variability.

## Main text

*SATB1* encodes a dimeric/tetrameric transcription factor^1^ with crucial roles in development and maturation of T-cells^2–4^. Recently, a potential contribution of *SATB1* to brain development was suggested by statistically significant enrichment of *de novo* variants in two large neurodevelopmental disorder (NDD) cohorts^5; 6^, although its function in the central nervous system is poorly characterized.

Through international collaborations^7–9^, we identified 42 individuals with a rare (likely) pathogenic variant in *SATB1* (NM_001131010.4), a gene under constraint against loss-of-function and missense variation (pLoF: o/e=0.15 (0.08-0.29); missense: o/e=0.46 (0.41-0.52); gnomAD v2.1.1)^10^. Twenty-eight of the *SATB1* variants occurred *de novo*, three were inherited from an affected parent, and five resulted from (suspected) parental mosaicism (Suppl. Figure 1). Reduced penetrance is suggested by two variants inherited from unaffected parents. Inheritance status of the final four could not be established (Suppl. Table 1A). Of note, two individuals carried a (likely) pathogenic variant in another known disease gene in addition to the *SATB1* variant, which (in part) explained the observed phenotype (*NF1*, MIM #162200 and *FOXP2*, MIM #602081). Thirty individuals carried 15 unique *SATB1* missense variants, including three recurrent variants (Figure 1A), significantly clustering in the highly homologous DNA-binding domains CUT1 and CUT2 (*p*=1.00e-7; Figure 2A, Suppl. Figure 2)^11; 12^. Ten individuals carried premature protein truncating variants (PTVs; two nonsense, seven frameshift, one splice site; Suppl. Table 1A, Suppl. Table 2), and two individuals had a (partial) gene deletion (Suppl. Figure 3). For 38 affected individuals and one mosaic parent, clinical information was available. Overall, we observed a broad phenotypic spectrum, characterized by neurodevelopmental delay (35/36, 97%), ID (28/31, 90%), muscle tone abnormalities (abnormal tone 28/37, 76%; hypotonia 28/37, 76%; spasticity 10/36, 28%), epilepsy (22/37, 61%) behavioral problems (24/34, 71%), facial dysmorphisms (24/36, 67%; Figure 1B-1D, Suppl. Figure 4A), and dental abnormalities (24/34, 71%) (Figure 1E, Table 1, Suppl. Figure 4B, Suppl. Table 1). Individuals with missense variants were globally more severely affected than those with PTVs: 57% of individuals with a missense variant had severe/profound ID whereas this level of ID was not observed for any individuals with PTVs. Furthermore, hypotonia, spasticity and (severe) epilepsy were more common in individuals with missense variants than in those with PTVs (92% versus 42%, 42% versus 0%, 80% versus 18%, respectively) (Figure 1G, Table 1, Suppl. Table 1A). To objectively quantify these observations, we divided our cohort into two variant-specific clusters (missense versus PTVs) and assessed the two groups using a Partitioning Around Medoids clustering algorithm^13^ on 100 features derived from standardized clinical data (Human Phenotype Ontology (HPO); Suppl. Figure 5A)^14^. Twenty-seven of the 38 individuals were classified correctly as either belonging to the PTV or missense variant group (*p*=0.022), confirming the existence of at least two separate clinical entities (Figure 1H, Suppl. Figure 5B). Moreover, computational averaging of facial photographs^15^ revealed clear differences between the average facial gestalt for individuals with missense variants when compared to individuals with PTVs or deletions (Figure 1B-F, Suppl. Figure 4, Suppl. Table 1B).

**Figure 1.**
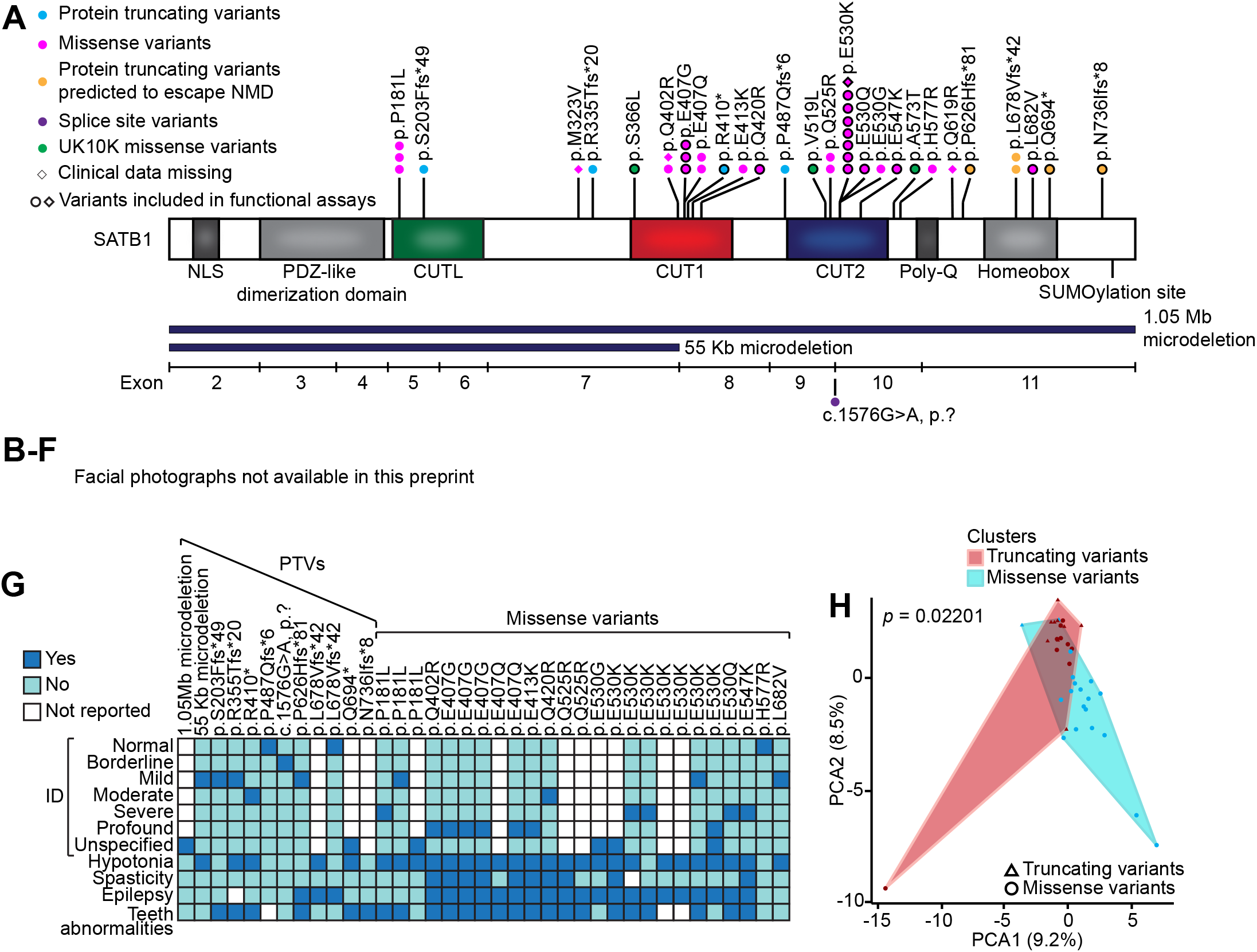
Clinical evaluation of *SATB1* variants in neurodevelopmental disorders. **A**) Schematic representation of the SATB1 protein (NM_001131010.4/NP_001124482.1), including functional domains, with truncating variants labeled in cyan, truncating variants predicted to escape NMD in orange, splice site variants in purple, missense variants in magenta, and UK10K rare control missense variants in green. Deletions are shown in dark blue below the protein schematic, above a diagram showing the exon boundaries. We obtained clinical data for all individuals depicted by a circle. **B-D**) Facial photographs of individuals with (partial) gene deletions and truncations predicted to result in haploinsufficiency (**B**), of individuals with truncations predicted to escape from NMD and resulting in transcriptionally active proteins (**C**) and of individuals with missense variants (**D**). All depicted individuals show facial dysmorphisms and although overlapping features are seen, no consistent facial phenotype can be observed for the group as a whole. Overlapping facial dysmorphisms include facial asymmetry, high forehead, prominent ears, straight and/or full eyebrows, puffy eyelids, downslant of palpebral fissures, low nasal bridge, full nasal tip and full nasal alae, full lips with absent cupid’s bow, prominent cupid’s bow or thin upper lip vermillion (Suppl. Table 1B). Individuals with missense variants are more alike than individuals in the truncating cohorts, and we observed recognizable overlap between several individuals in the missense cohort (individual 17, 27, 31, 37, the siblings 19, 20 and 21, and to a lesser extent individual 24 and 35). A recognizable facial overlap between individuals with the other two variant types could not be observed. Related individuals are marked with a blue box. **E**) Photographs of teeth abnormalities observed in individuals with *SATB1* variants. Dental abnormalities are seen for all variant types and include widely spaced teeth, dental fragility, missing teeth, disorganized teeth implant, and enamel discoloration (Suppl. Table 1B). **F**) Computational average of facial photographs of 16 individuals with a missense variant (left) and 8 individuals with PTVs or (partial) gene deletions (right). **G**) Mosaic plot presenting a selection of clinical features. Individuals with no or very limited clinical data were omitted. **H**) The Partitioning Around Medoids analysis of clustered HPO-standardized clinical data from 38 individuals with truncating (triangle) and missense variants (circle) shows a significant distinction between the clusters of individuals with missense variants (blue) and individuals with PTVs (red). Applying Bonferroni correction, a *p*-value smaller than 0.025 was considered significant.

**Figure 2.**
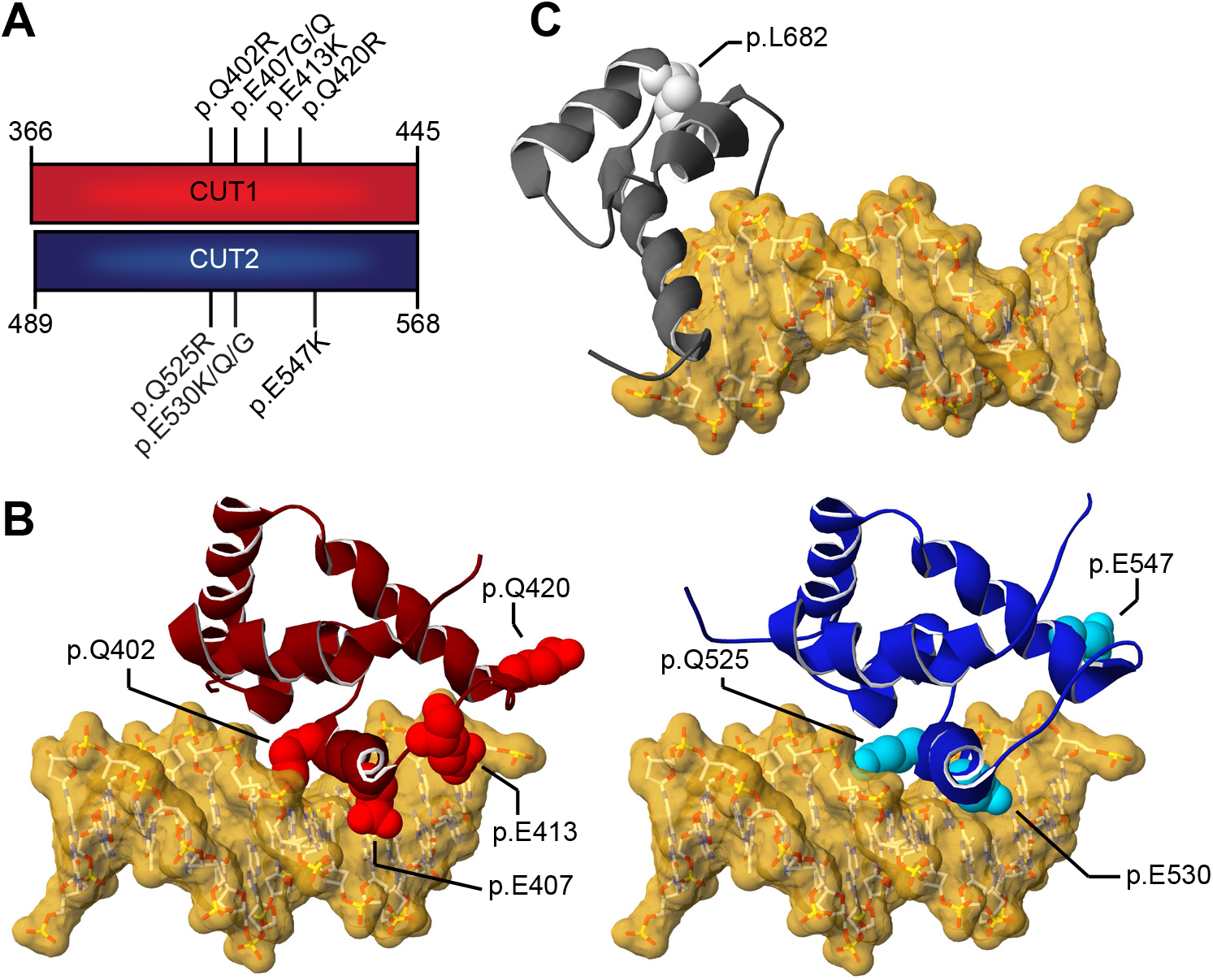
3D protein modeling of SATB1 missense variants in DNA-binding domains. **A**) Schematic representation of the aligned CUT1 and CUT2 DNA-binding domains. CUT1 and CUT2 domains have a high sequence identity (40%) and similarity (78%). Note that the recurrent p.Q402R, p.E407G/p.E407Q and p.Q525R, p.E530G/p.E530K/p.E530Q variants affect equivalent positions within the respective CUT1 and CUT2 domains, while p.Q420R in CUT1 and p.E547K in CUT2 affect cognate regions. **B**) 3D-model of the SATB1 CUT1 domain (left; PDB 2O4A) and CUT2 domain (right; based on PDB 2CSF) in interaction with DNA (yellow). Mutated residues are highlighted in red for CUT1 and cyan for CUT2, along the ribbon visualization of the corresponding domains in burgundy and dark blue, respectively. **C**) 3Dhomology model of the SATB1 homeobox domain (based on PDB 1WI3 and 2D5V) in interaction with DNA (yellow). The mutated residue is shown in light gray along the ribbon visualization of the corresponding domain in dark gray. **B-C**) For more detailed descriptions of the different missense variants in our cohort, see Suppl. Notes.

**Table 1.** Summary of clinical characteristics associated with (de novo) *SATB1* variants

|  | All individuals |  | Individuals with PTVs and (partial) gene deletions |  | Individuals with missense variants |  |
| --- | --- | --- | --- | --- | --- | --- |
|  | % | Present / total assessed | % | Present / total assessed | % | Present / total assessed |
| **Neurologic** |  |  |  |  |  |  |
| Intellectual disability | 90 | 28/31 | 80 | 8/10 | 95 | 20/21 |
| Normal | 10 | 3/31 | 20 | 2/10 | 5 | 1/21 |
| Borderline | 0 | 0/31 | 0 | 0/10 | 0 | 0/21 |
| Mild | 26 | 8/31 | 60 | 6/10 | 10 | 2/21 |
| Moderate | 10 | 3/31 | 10 | 1/10 | 10 | 2/21 |
| Severe | 19 | 6/31 | 0 | 0/10 | 29 | 6/21 |
| Profound | 19 | 6/31 | 0 | 0/10 | 29 | 6/21 |
| Unspecified | 16 | 5/31 | 10 | 1/10 | 19 | 4/21 |
| Developmental delay | 97 | 35/36 | 100 | 12/12 | 96 | 23/24 |
| Motor delay | 92 | 34/37 | 92 | 11/12 | 92 | 23/25 |
| Speech delay | 89 | 32/36 | 83 | 10/12 | 92 | 22/24 |
| Dysarthria | 30 | 6/20 | 9 | 1/11 | 56 | 5/9 |
| Epilepsy | 61 | 22/36 | 18 | 2/11 | 80 | 20/25 |
| EEG abnormalities | 79 | 19/24 | 29 | 2/7 | 100 | 17/17 |
| Hypotonia | 76 | 28/37 | 42 | 5/12 | 92 | 23/25 |
| Spasticity | 28 | 10/36 | 0 | 0/12 | 42 | 10/24 |
| Ataxia | 22 | 6/27 | 17 | 2/12 | 27 | 4/15 |
| Behavioral disturbances | 71 | 24/34 | 58 | 7/12 | 77 | 17/22 |
| Sleep disturbances | 41 | 12/29 | 27 | 3/11 | 50 | 9/18 |
| Abnormal brain imaging | 55 | 17/31 | 43 | 3/7 | 58 | 14/24 |
| Regression | 17 | 6/35 | 8 | 1/12 | 22 | 5/23 |
| **Growth** |  |  |  |  |  |  |
| Abnormalities during pregnancy | 24 | 8/33 | 27 | 3/11 | 23 | 5/22 |
| Abnormalities during delivery | 32 | 10/31 | 55 | 6/11 | 20 | 4/20 |
| Abnormal term of delivery | 6 | 2/31 | 10 | 1/10 | 5 | 1/21 |
| Preterm (<37 weeks) | 6 | 2/31 | 10 | 1/10 | 5 | 1/21 |
| Postterm (>42 weeks) | 0 | 0/31 | 0 | 0/10 | 0 | 0/21 |
| Abnormal weight at birth | 16 | 5/32 | 22 | 2/9 | 13 | 3/23 |
| Small for gestational age (<p10) | 9 | 3/32 | 11 | 1/9 | 9 | 2/23 |
| Large for gestational age (>p90) | 6 | 2/32 | 11 | 1/9 | 4 | 1/23 |
| Abnormal head circumference at birth | 7 | 1/14 | 17 | 1/6 | 0 | 0/8 |
| Microcephaly* | 0 | 0/14 | 0 | 0/6 | 0 | 0/8 |
| Macrocephaly# | 7 | 1/14 | 17 | 1/6 | 0 | 0/8 |
| Abnormal height | 21 | 6/29 | 9 | 1/11 | 28 | 5/18 |
| Short stature* | 14 | 4/29 | 0 | 0/11 | 22 | 4/18 |
| Tall stature# | 7 | 2/29 | 9 | 1/11 | 6 | 1/18 |
| Abnormal head circumference | 26 | 7/31 | 11 | 1/9 | 32 | 6/22 |
| Microcephaly* | 26 | 7/31 | 11 | 1/9 | 32 | 6/22 |
| Macrocephaly# | 0 | 0/31 | 0 | 0/9 | 0 | 0/22 |
| Abnormal weight | 48 | 13/27 | 11 | 1/9 | 67 | 12/18 |
| Underweight* | 22 | 6/27 | 11 | 1/9 | 28 | 5/18 |
| Overweight# | 26 | 7/27 | 0 | 0/9 | 39 | 7/18 |
| **Other phenotypic features** |  |  |  |  |  |  |
| Facial dysmorphisms | 67 | 24/36 | 64 | 7/11 | 68 | 17/25 |
| Dental/oral abnormalities | 71 | 24/34 | 55 | 6/11 | 78 | 18/23 |
| Drooling/dysphagia | 38 | 12/32 | 25 | 3/12 | 45 | 9/20 |
| Hearing abnormalities | 7 | 2/30 | 18 | 2/11 | 0 | 0/19 |
| Vision abnormalities | 55 | 17/31 | 73 | 8/11 | 45 | 9/20 |
| Cardiac abnormalities | 19 | 6/32 | 27 | 3/11 | 14 | 3/21 |
| Skeleton/limb abnormalities | 38 | 13/34 | 18 | 2/11 | 48 | 11/23 |
| Hypermobility of joints | 30 | 8/27 | 30 | 3/10 | 29 | 5/17 |
| Gastrointestinal abnormalities | 53 | 17/32 | 27 | 3/11 | 67 | 14/21 |
| Urogenital abnormalities | 17 | 5/30 | 0 | 0/11 | 26 | 5/19 |
| Endocrine/metabolic abnormalities | 30 | 9/30 | 0 | 0/11 | 47 | 9/19 |
| Immunological abnormalities | 32 | 8/25 | 25 | 2/8 | 35 | 6/17 |
| Skin/hair/nail abnormalities | 24 | 8/34 | 9 | 1/11 | 30 | 7/23 |
| Neoplasms in medical history | 0 | 0/34 | 0 | 0/11 | 0 | 0/23 |

\* <p3

We performed functional analyses assessing consequences of different types of *SATB1* variants for cellular localization, transcriptional activity, overall chromatin binding, and dimerization capacity. Based on protein modeling (Figure 2, Suppl. Notes), we selected five missense variants in CUT1 and CUT2 affecting residues that interact with, or are close to, the DNA backbone (mosaic variant p.E407G and *de novo* variants p.Q420R, p.E530K/p.E530Q, p.E547K), as well as the only homeobox domain variant (p.L682V, *de novo*). As controls, we selected three rare missense variants from the UK10K consortium, identified in healthy individuals with a normal IQ: p.S366L (gnomAD allele frequency 6.61e-4), p.V519L (8.67e-6) and p.A573T (1.17e-4) (Figure 1A, Suppl. Table 3)^16^. When overexpressed as YFP-fusion proteins in HEK293T/17 cells, wildtype SATB1 localized to the nucleus in a granular pattern, with an intensity profile inverse to the DNA-binding dye Hoechst 33342 (Figure 3A-B). In contrast to wildtype and UK10K control missense variants, the p.E407G, p.Q420R, p.E530K/p.E530Q and p.E547K variants displayed a cage-like clustered nuclear pattern, strongly co-localizing with the DNA (Figure 3A-B, Suppl. Figure 6).

**Figure 3.**
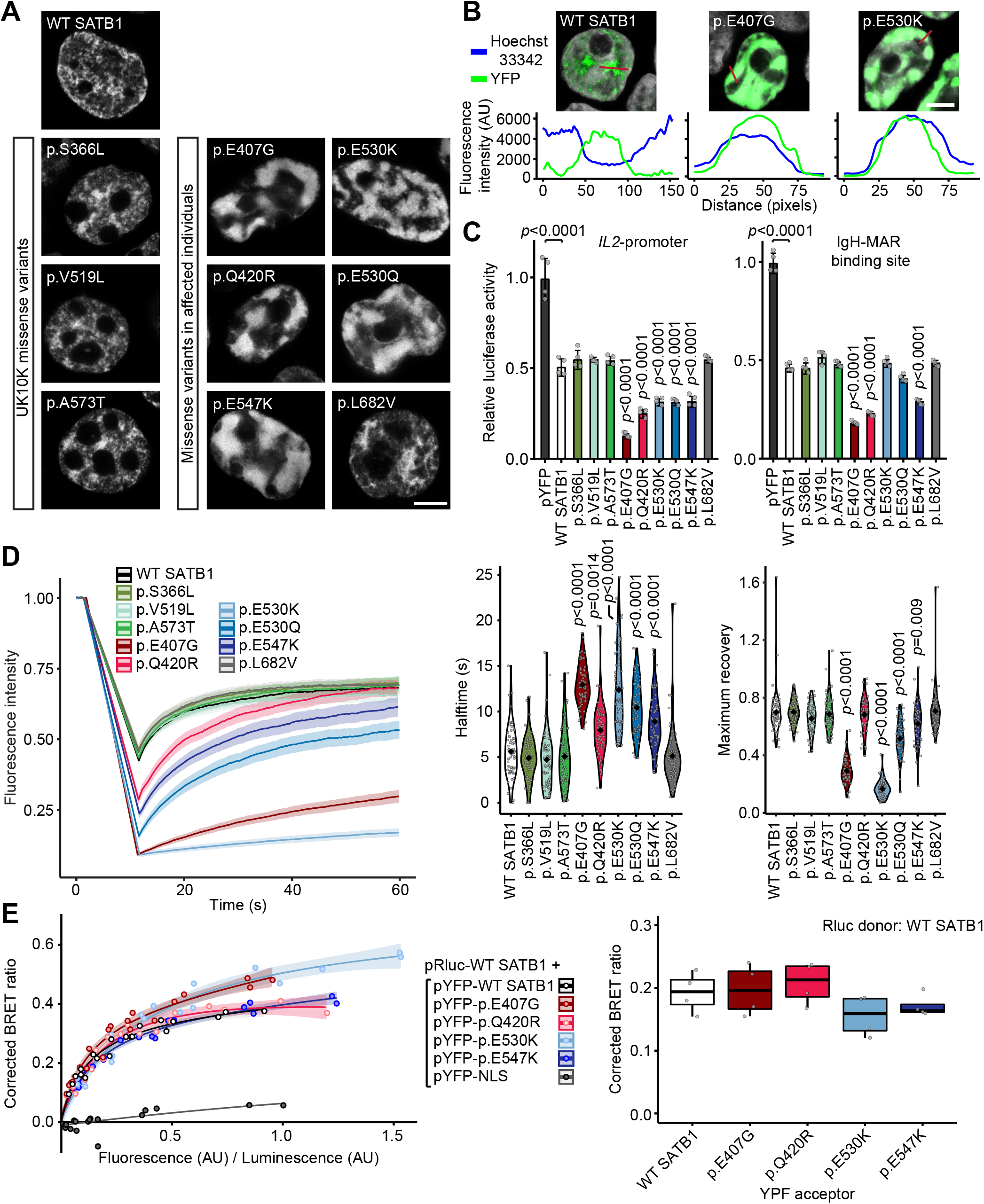
SATB1 missense variants stabilize DNA binding and show increased transcriptional repression. **A**) Direct fluorescence super-resolution imaging of nuclei of HEK293T/17 cells expressing YFP-SATB1 fusion proteins. Scale bar=5μm. **B**) Intensity profiles of YFP-tagged SATB1 and variants, and the DNA binding dye Hoechst 33342. The graphs represent the fluorescence intensity values of the position of the red lines drawn in the micrographs on the top (SATB1 proteins in green, Hoechst 33342 in white, scale bar = 5 μm). For each condition a representative image and corresponding intensity profile plot is shown. **C**) Luciferase reporter assays using reporter constructs containing the *IL2*-promoter region and the IgH matrix associated region (MAR) binding site. UK10K control variants are shaded in green, CUT1 domain variants in red, CUT2 domain variants in blue and the homeobox variant in gray. Values are expressed relative to the control (pYFP; black) and represent the mean ± S.E.M. (*n* = 4, *p*-values compared to wildtype SATB1 (WT; white), one-way ANOVA and *post-hoc* Bonferroni test). **D**) FRAP experiments to assess the dynamics of SATB1 chromatin binding in live cells. Left, mean recovery curves ± 95% C.I. recorded in HEK293T/17 cells expressing YFP-SATB1 fusion proteins. Right, violin plots with median of the halftime (central panel) and maximum recovery values (right panel) based on single-term exponential curve fitting of individual recordings (*n* = 60 nuclei from three independent experiments, *p*-values compared to WT SATB1, one-way ANOVA and *post-hoc* Bonferroni test). Color code as in C. **E**) BRET assays for SATB1 dimerization in live cells. Left, mean BRET saturation curves ± 95% C.I. fitted using a non-linear regression equation assuming a single binding site (*y* = BRETmax * *x* / (BRET50 / *x*); GraphPad). The corrected BRET ratio is plotted against the ratio of fluorescence/luminescence (AU) to correct for expression level differences between conditions. Right, corrected BRET ratio values at mean BRET50 level of WT SATB1, based on curve fitting of individual experiments (*n* = 4, one-way ANOVA and *post-hoc* Bonferroni test, no significant differences). Color code as in C. **A-E**) When compared to WT YFP-SATB1 or UK10K variants, most variants identified in affected individuals show a nuclear cage-like localization (**A**), stronger co-localization with the DNA-binding dye Hoechst 33342 (**B**), increased transcriptional repression (**C**), reduced protein mobility (**D**) and unchanged capacity of interaction with WT SATB1 (**E**).

To assess the effects of SATB1 missense variants on transrepressive activity, we used a luciferase reporter system with two previously established downstream targets of SATB1, the *IL2*-promoter and IgH-MAR (matrix associated region)^17–19^. All five functionally assessed CUT1 and CUT2 missense variants demonstrated increased transcriptional repression of the *IL2*-promoter, while the UK10K control variants did not differ from wildtype (Figure 3C). In assays using IgH-MAR, increased repression was seen for both CUT1 variants, and for one of the CUT2 variants (Figure 3C). Taken together, these data suggest that etiological SATB1 missense variants in CUT1 and CUT2 lead to stronger binding of the transcription factor to its targets.

To study whether SATB1 missense variants affect the dynamics of chromatin binding more globally, we employed fluorescent recovery after photobleaching (FRAP) assays. Consistent with the luciferase reporter assays, all CUT1 and CUT2 missense variants, but not the UK10K control variants, affected protein mobility in the nucleus. The CUT1 and CUT2 variants demonstrated increased halftimes and reduced maximum recovery, suggesting stabilization of SATB1 chromatin binding (Figure 3D).

In contrast to the CUT1 and CUT2 missense variants, the homeobox variant p.L682V did not show functional differences from wildtype (Figure 3A-D, Suppl. Figure 6), suggesting that, although it is absent from gnomAD, highly intolerant to variation and evolutionarily conserved (Suppl. Figure 2, Suppl. Figure 7A-B), this variant is unlikely to be pathogenic. This conclusion is further supported by the presence of a valine residue at the equivalent position in multiple homologous homeobox domains (Suppl. Figure 7C). Additionally, the mild phenotypic features in this individual (individual 42) can be fully explained by an out-of-frame *de novo* intragenic duplication of *FOXP2*, known to cause disease through haploinsufficiency^20^.

We went on to assess the impact of the CUT1 and CUT2 missense variants (p.E407G, p.Q420R, p.E530K, p.E547K) on protein interaction capacities using bioluminescence resonance energy transfer (BRET). All tested variants retained the ability to interact with wildtype SATB1 (Figure 3E), with the potential to yield dominant-negative dimers/tetramers *in vivo* and to disturb normal activity of the wildtype protein.

The identification of *SATB1* deletions suggests that haploinsufficiency is a second underlying disease mechanism. This is supported by the constraint of *SATB1* against loss-of-function variation, and the identification of PTV carriers that are clinically distinct from individuals with missense variants. PTVs are found throughout the locus and several are predicted to undergo NMD by *in silico* models of NMD efficacy (Suppl. Table 4)^21^. In contrast to these predictions, we found that one of the PTVs, p.R410*, escapes NMD (Suppl. Figure 8A-B). However, the p.R410* variant would lack critical functional domains (CUT1, CUT2, homeobox) and indeed showed reduced transcriptional activity in luciferase reporter assays when compared to wildtype protein (Suppl. Figure 8), consistent with the haploinsufficiency model.

Four unique PTVs that we identified were located within the final exon of SATB1 (Figure 1A) and predicted to escape NMD (Suppl. Table 4). Following experimental validation of NMD escape (Figure 4A-B), three such variants (p.P626Hfs*81, p.Q694* and p.N736Ifs*8) were assessed with the same functional assays that we used for missense variants. When overexpressed as YFP-fusion proteins, the tested variants showed altered subcellular localization, forming nuclear puncta or (nuclear) aggregates, different from patterns observed for missense variants (Figure 4C, Suppl. Figure 9A-B). In luciferase reporter assays, the p.P626Hfs*81 variant showed increased repression of both the *IL2*-promoter and IgH-MAR, whereas p.Q694* only showed reduced repression of IgH-MAR (Figure 4D). The p.N736Ifs*8 variant showed repression comparable to that of wildtype protein for both targets (Figure 4D). In further pursuit of pathophysiological mechanisms, we tested protein stability and SUMOylation, as the previously described p.K744 SUMOylation site is missing in all assessed NMD-escaping truncated proteins (Figure 4A)^22^. Our observations suggest the existence of multiple SATB1 SUMOylation sites (Suppl. Figure 10) and no effect of NMD-escaping variants on SUMOylation of the encoded proteins (Suppl. Figure 10) nor any changes in protein stability (Suppl. Figure 9C). Although functional assays with NMD-escaping PTVs hint towards additional disease mechanisms, HPO-based phenotypic analysis could not confirm a third distinct clinical entity (p=0.932; Suppl. Figure 4E, Suppl. Table 5).

**Figure 4.**
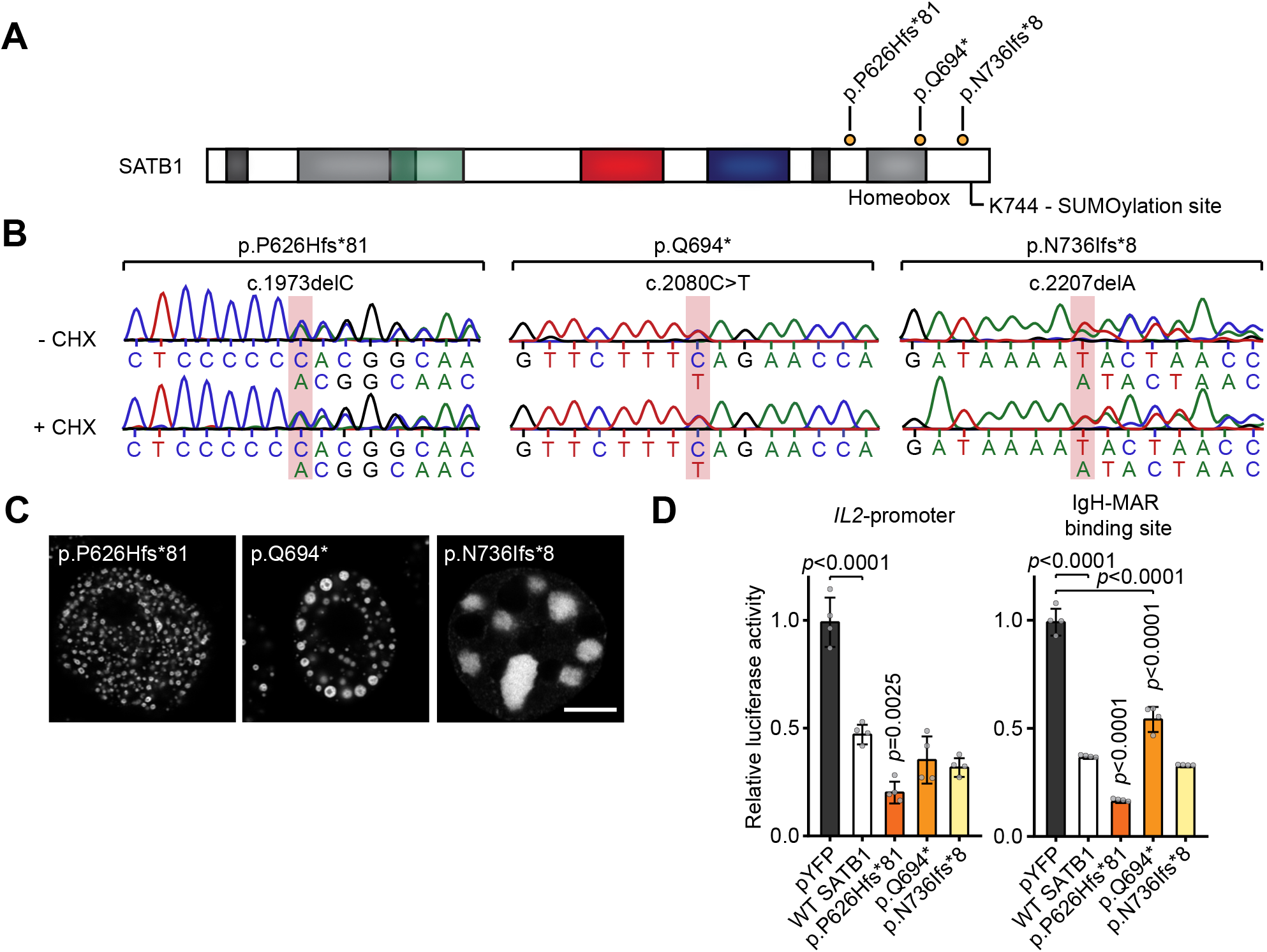
*SATB1* frameshift variants in the last exon escape NMD. **A**) Schematic overview of the SATB1 protein, with truncating variants predicted to escape NMD that are included in functional assays labeled in orange. A potential SUMOylation site at position p.K744 is highlighted. **B**) Sanger sequencing traces of patient-derived EBV immortalized lymphoblastoid cell lines treated with or without cycloheximide (CHX) to test for NMD. The mutated nucleotides are shaded in red. Transcripts from both alleles are present in both conditions showing that these variants escape NMD. **C**) Direct fluorescence super-resolution imaging of nuclei of HEK293T/17 cells expressing SATB1 truncating variants fused with a YFP-tag. Scale bar = 5 μm. Compared to WT YFP-SATB1, NMD-escaping variants show altered localization forming nuclear puncta or aggregates. **D**) Luciferase reporter assays using reporter constructs containing the *Il2*-promoter and the IgH matrix associated region (MAR) binding site. Values are expressed relative to the control (pYFP; black) and represent the mean ± S.E.M. (*n* = 4, *p*-values compared to WT SATB1 (white), one-way ANOVA and post-hoc Bonferroni test). All NMD-escaping variants are transcriptionally active and show repression of the *Il2*-promoter and IgH-MAR binding site.

Our study demonstrates that while statistical analyses^5; 6^ can provide the first step towards identification of new NDDs, a mutation-specific functional follow-up is required to gain insight into the underlying mechanisms and to understand phenotypic differences within patient cohorts. Multiple mechanisms and/or more complex genotype-phenotype correlations are increasingly appreciated in newly described NDDs, such as those associated with *RAC1, POL2RA, KMT2E* and *PPP2CA*^23–26^. Interestingly, although less often explored, such mechanistic complexity might also underlie well-known (clinically recognizable) NDDs. For instance, a CUT1 missense variant in *SATB2*, a paralog of *SATB1* that causes Glass syndrome through haploinsufficiency (MIM #612313)^27^, affects protein localization and nuclear mobility in a similar manner as the corresponding *SATB1* missense variants (Suppl. Figure 11, Suppl. Figure 12)^28^. Taken together, these observations suggest that mutation-specific mechanisms await discovery both for new and well-established clinical syndromes.

In summary, we demonstrate that at least two different previously uncharacterized NDDs are caused by distinct classes of rare (*de novo*) variation at a single locus. We combined clinical investigation, *in silico* models and cellular assays to characterize the phenotypic consequences and functional impacts of a large patient series uncovering distinct pathophysiological mechanisms of the *SATB1*-associated NDDs. This level of combined analyses is recommended for known and yet undiscovered NDDs to fully understand disease etiology.

## Acknowledgements

We are extremely grateful to all families participating in this study. In addition, we wish to thank the members of the Genome Technology Center and Cell culture facility, Department of Human Genetics, Radboud university medical center, Nijmegen, for data processing and cell culture of patient-derived cell lines. This work was financially supported by Aspasia grants of the Dutch Research Council (015.014.036 to TK and 015.014.066 to LV), Netherlands Organization for Health Research and Development (91718310 to TK), the Max Planck Society (JdH, SF), Oxford Brookes University, the Leverhulme Trust and the British Academy (DN), and grants from the Swiss National Science Foundation (31003A_182632 to AR), Lithuanian-Swiss cooperation program to reduce economic and social disparities within the enlarged European Union (AR, VK) and the Jérôme Lejeune Foundation (AR). In individual 13, 14 and 15, whole-exome sequencing was performed in the framework of the German project “TRANSLATE NAMSE”, an initiative from the National Action League for People with Rare Diseases (Nationales Aktionsbündnis für Menschen mit Seltenen Erkrankungen, NAMSE) facilitating innovative genetic diagnostics for individuals with suggested rare diseases. Part of this work (IT) was performed under the Genomic Answers for Kids program funded by generous donors to the Children’s Mercy Research Institute. We wish to thank all the ALSPAC families who took part in this study, the midwives for their help in recruiting them, and the whole ALSPAC team, which includes interviewers, computer and laboratory technicians, clerical workers, research scientists, volunteers, managers, receptionists and nurses. The UK Medical Research Council and Wellcome (Grant ref: 102215/2/13/2) and the University of Bristol provide core support for ALSPAC. A comprehensive list of grants funding is available on the ALSPAC website (http://www.bristol.ac.uk/alspac/external/documents/grant-acknowledgements.pdf). Wh ole-genome sequencing of the ALSPAC samples was performed as part of the UK10K consortium (a full list of investigators who contributed to the generation of the data is available from www.UK10K.org.uk). This research was made possible through access to the data and findings generated by the 100,000 Genomes Project. The 100,000 Genomes Project is managed by Genomics England Limited (a wholly owned company of the Department of Health and Social Care). The 100,000 Genomes Project is funded by the National Institute for Health Research and NHS England. The Wellcome Trust, Cancer Research UK and the Medical Research Council have also funded research infrastructure. The 100,000 Genomes Project uses data provided by patients and collected by the National Health Service as part of their care and support. In addition, the collaborations in this study were facilitated by ERN ITHACA, one of the 24 European Reference Networks (ERNs) approved by the ERN Board of Member States, co-funded by European Commission. For more information about the ERNs and the EU health strategy please visit http://ec.europa.eu/health/ern. The aims of this study contribute to the Solve-RD project (EdB, HB, CG, AJ, TK, LV) which has received funding from the European Union’s Horizon 2020 research and innovation programme under grant agreement No 779257.

## Conflict of interest

KMc, TBP, and TSS are employees of GeneDx, Inc. KR is employee of Ambrygen Genetics.

## Contributions

JdH, EdB, AR, SF and LV designed the study. Clinical data collection and interpretation was performed by EdB and TK. AD and JHK carried out HPO-based clustering analysis. Facial averaging was done by CN. The three-dimensional modeling was performed by NG. LW and CG did the mutational clustering analysis. JdH designed and executed cell-based assays. DN suggested use of healthy UK10K controls for functional assays and provided accompanying phenotypic data. EdB, NV, SA, SB, FB, BBZ, VB, ALB, TB, HB, HC, JC, LC, HCo, DDD, ED, FD, ASDP, CD, DD, DDy, OE, LF, LG, BH, YH, YHA, JH, BH, TI, AJ, RJ, KJ, SJ, HK, MK, AKM, UK, FK, UKo, VK, VKu, AK, AL, PL, LM, NM, BM, KMc, VM, MM, SM, TM, LMo, SMo, HMS, HM, DN, RNE, SO, LO, MO, TBP, MP, AP, RP, EP, KR, ER, JR, TSS, CS, MS, LSB, RS, AS, IT, LT, AVD, JVdS, SV, EV, AV, MW, AW, MWi, BZ, TK participated in recruitment of individuals, phenotyping and/or next-generation sequencing analysis. JdH, EdB, AR, SF and LV analyzed and interpreted the results and wrote the manuscript. AR, SF and LV supervised the project. All authors contributed to the final version of the manuscript.

## Methods

### Individuals and consent

For all individuals reported in this study, informed consent was obtained to publish unidentifiable data. When applicable, specific consent was obtained for publication of clinical photographs and inclusion of photographs in facial analysis. All consent procedures are in accordance with both the local ethical guidelines of the participating centers, and the Declaration of Helsinki. Individuals with possible (likely) pathogenic *SATB1* variants were identified through international collaborations facilitated by MatchMakerExchange^7^, GPAP of RD-connect^8^, the Solve-RD consortium, the Decipher Database^9^, and through searching literature for cohort-studies for NDD^5; 6^. Individuals 27 and 28 were previously described in a clinical case report^29^. Clinical characterization was performed by reviewing the medical files and/or revising the phenotype of the individuals in the clinic. All (affected) individuals with a *SATB1* variant are included in Suppl. Table 1. A summary of clinical characteristics is provided in Table 1, including 38 of 42 individuals: individual 16, 32 and 41 were excluded because no clinical data were available, individual 22 was excluded as she is (low) mosaic for the *SATB1* variant (~1%). In Figure 1G, 37 of 42 individuals were included: in addition to individuals 16, 22, 32, and 41, we also excluded individual 18, for whom only very limited clinical information was available.

### Next generation sequencing

For all individuals except individual 1, 2, and 28, *SATB1* variants were identified by whole exome sequencing after variant filtering as previously described^11; 30–35^. Information on inheritance was obtained after parental confirmation, either from parental exome sequencing data or through targeted Sanger sequencing. For individual 1 the *SATB1* variant was identified by array-CGH and for individual 2 an Affymetrix Cytoscan HD array was performed in addition to whole exome sequencing. For individual 28 targeted Sanger sequencing was performed after identification of the variant in his similarly affected sister. To predict deleteriousness of variants, CADD-PHRED V1.4 scores and SpliceAI scores (VCFv4.2; dated 20191004) were obtained for all variants identified in affected individuals^36; 37^. In addition, for all nonsense, frameshift and splice site variants, NMDetective scores were obtained (v2)^21^. For all missense variants, we analysed the mutation tolerance of the site of the affected residue using Metadome^38^.

### UK10K controls for functional assays

Genome sequence data from 1,867 ALSPAC^39; 40^ individuals in the UK10K^16^ dataset were annotated in ANNOVAR^41^ and filtered to identify individuals carrying rare coding variants (gnomAD genome_ALL frequency<0.1%) within *SATB1*. In total six rare variants were identified. These variants were carried by 13 individuals, all in a heterozygous state. Three variants (one in the CUT1 domain, one in the CUT2 domain and one outside of critical domains) were selected for functional studies. These variants were carried by nine individuals. Phenotypic data of carriers and non-carriers were available through the ALSPAC cohort, an epidemiological study of pregnant women who were resident in Avon, UK with expected dates of delivery 1st April 1991 to 31st December 1992. This dataset included 13,988 children who were alive at 1 year of age, 1,867 of whom underwent genome sequencing as part of the UK10K project. Of the UK10K individuals, 1,741 children had measures of IQ (WISC) collected at age 8 years providing an indication of cognitive development. The ALSPAC study website contains details of all the data that is available through a fully searchable data dictionary and variable search tool (http://www.bristol.ac.uk/alspac/researchers/our-data/)

### Human Phenotype Ontology (HPO)-based phenotype clustering analysis

All clinical data were standardized using HPO terminology^14^. Thirty-eight of 42 individuals were included in analysis: individual 16, 32 and 41 were excluded because no clinical data were available, individual 22 was excluded as she is (low) mosaic for the SATB1 variant (~1%). The semantic similarity between all the HPO terms used in this cohort (356 features) was calculated using the Wang algorithm in the HPOSim package^42; 43^ in R. HPO terms with at least a 0.5 similarity score were grouped (Suppl. Figure 5): a new feature was created as a replacement, which was the sum of the grouped features. For eleven terms, the HPO semantic similarity could not be calculated using HPOSim. Seven of those could be manually assigned to a group, since the feature clearly matched (for instance: nocturnal seizures with the seizure/epilepsy group). For a full list of the grouped features, see Suppl. Table 6. HPO terms that could not be grouped were added as separate features, as was severity of intellectual disability. This led to 100 features for every individual, instead of the previous 356 separate HPO terms. To quantify the possible genotype/phenotype correlation in the cohort, we used Partitioning Around Medoids (PAM) clustering^13^ dividing our cohort into two groups (missense variants versus truncating variants), followed by a permutations test (*n*=100,000) and relabeling based on variant types, while keeping the original distribution of variant types into account. The same clustering and permutations test was performed when dividing our cohort into three groups. For both analyses, Bonferroni correction for multiple testing was applied and a *p*-value smaller than 0.025 was considered significant.

### Average face analysis

For 24 of 42 individuals facial 2D-photographs were available for facial analysis. As previously described, average faces were generated while allowing for asymmetry preservation and equal representation by individuals^15^.

### Three-dimensional protein modeling

The crystal structure of the CUT1 domain of SATB1 bound to Matrix Attachment Region DNA (PDB entry 2O4A^44^) was used to contextualize the SATB1 CUT1 variants with respect to DNA using Swiss-PdbViewer^45^. The solution structure of the CUT2 domain of human SATB2 (first NMR model of the PDB entry 2CSF^46^) was used as a template to align the SATB1 residues T491 to H577 (Uniprot entry Q01826), and to build a model using Swiss-PdbViewer^45^. The model of the CUT2 domain was superposed onto the SATB1 CUT1 domain bound to Matrix Attachment Region DNA (PDB entry 2O4A^44^ using the “magic fit” option of Swiss-PdbViewer^45^) to contextualize the SATB1 CUT2 variants with respect to DNA. The solution structure of the homeodomain of human SATB2 (second NMR model of the PDB entry 1WI3^47^ was used as a template to align SATB1 residues P647 to G704 (Uniprot entry Q01826), and to build a model using Swiss-PdbViewer^45^. Chains A, C and D of the crystal structure of HNF-6alpha DNA-binding domain in complex with the TTR promoter (PDB entry 2D5V), which has a DNA binding domain similar to the CUT2 domain of SATB1 and a second DNA binding domain similar to the homeobox of SATB1, was used as a template to superpose the model of the SATB2 homeobox domain onto the HNF-6alpha structure using the “magic fit” option of Swiss-PdbViewer^48^ to contextualize the SATB1 homeobox variant with respect to DNA.

### Spatial clustering analysis of missense variants

Twenty-four of the observed 30 missense variants were included in the spatial clustering analysis. We excluded 6 variants, to correct for familial occurrence. The geometric mean was computed over the locations of observed (*de novo*) missense variants in the cDNA of *SATB1* (NM_001131010.4). This geometric mean was then compared to 1,000,000 permutations, by redistributing the (*de novo*) variant locations over the total size of the coding region of *SATB1* (2,388 bp) and calculating the resulting geometric mean from each of these permutations. The *p*-value was then computed by checking how often the observed geometric mean distance was smaller than the permutated geometric mean distance. This approach was previously used to identify cDNA clusters of variants^11; 12^. Code used in this analysis is available at: https://github.com/laurensvdwiel/SpatialClustering.

### DNA expression constructs and site-directed mutagenesis

The cloning of SATB1 (NM_001131010.4), SATB2 (NM_001172509) and SUMO1 (NM_003352.4), has been described previously^49; 50^. Variants in SATB1 and SATB2 were generated using the QuikChange Lightning Site-Directed Mutagenesis Kit (Agilent). The primers used for site-directed mutagenesis are listed in Suppl. Table 7. cDNAs were subcloned using *BamHI/XbaI* (SATB1 and SUMO1) and *BclI/XbaI* (SATB2) restriction sites into pRluc and pYFP, created by modification of the pEGFP-C2 vector (Clontech) as described before^51^. To generate a UBC9-SATB1 fusion, the UBC9 (NM_194260.2) and SATB1 coding sequences were amplified using primers listed in Suppl. Table 8, and subcloned into the pHisV5 vector (a modified pEGFP-C2 vector adding an N-terminal His-and V5-tag) using *BamHI/SmaI* (UBC9) and *HindIII/XhoI* (SATB1) restriction sites. All constructs were verified by Sanger sequencing.

### Cell culture

HEK293T/17 cells (CRL-11268, ATCC) were cultured in DMEM supplemented with 10% fetal bovine serum and 1x penicillin-streptomycin (all Invitrogen) at 37°C with 5% CO_2_. Transfections for functional assays were performed using GeneJuice (Millipore) following the manufacturer’s protocol. Lympoblastoid cell lines (LCLs) were established by Epstein-Barr virus transformation of peripheral lymphocytes from blood samples collected in heparin tubes, and maintained in RPMI medium (Sigma) supplemented with 15% fetal bovine serum and 5% HEPES (both Invitrogen).

### Testing for nonsense mediated decay of truncating variants

Patient-derived LCLs were grown for 4 h with 100 μg/ml cycloheximide (Sigma) to block NMD. After treatment, cell pellets (10*10^6^ cells) were collected and RNA was extracted using the RNeasy Mini Kit (Qiagen). RT-PCR was performed using SuperScriptIII Reverse Transcriptase (ThermoFisher) with random primers, and regions of interest were amplified from cDNA using primers listed in Suppl. Table 9.

### Fluorescence microscopy

HEK293T/17 cells were grown on coverslips coated with poly-D-lysine (Sigma). Cells were fixed with 4% paraformaldehyde (PFA, Electron Microscopy Sciences) 48 h after transfection with YFP-tagged SATB1 and SATB2 variants. Nuclei were stained with Hoechst 33342 (Invitrogen). Fluorescence images were acquired with a Zeiss LSM880 confocal microscope and ZEN Image Software (Zeiss). For images of single nuclei, the Airyscan unit (Zeiss) was used with a 4.5 zoom factor. All other images were acquired with a 2.0 zoom factor. Intensity profiles were plotted using the ‘Plot Profile’ tool in Fiji - ImageJ.

### FRAP assays

HEK293T/17 cells were transfected in clear-bottomed black 96-well plates with YFP-tagged SATB1 and SATB2 variants. After 48 h, medium was replaced with phenol red-free DMEM supplemented with 10% fetal bovine serum (both Invitrogen), and cells were moved to a temperature-controlled incubation chamber at 37°C. Fluorescent recordings were acquired using a Zeiss LSM880 and Zen Black Image Software, with an alpha Plan-Apochromat 100x/1.46 Oil DIC M27 objective (Zeiss). FRAP experiments were performed by photobleaching an area of 0.98 μm x 0.98 μm within a single nucleus with 488-nm light at 100% laser power for 15 iterations with a pixel dwell time of 32.97 μs, followed by collection of times series of 150 images with a 2.5 zoom factor and an optical section thickness of 1.4 μm (2.0 Airy units). Individual recovery curves were background subtracted and normalized to the pre-bleach values, and mean recovery curves were calculated using EasyFRAP software^52^. Curve fitting was done with the FrapBot application using direct normalization and a single-component exponential model, to calculate the half-time and maximum recovery^53^.

### Luciferase reporter assays

Luciferase reporter assays were performed with a pIL2-luc reporter construct containing the human *IL2*-promoter region, and a pGL3-basic firefly luciferase reporter plasmid carrying seven repeats of the-TCTTTAATTTCTAATATATTTAGAAttc-MAR sequence identified in an enhancer region 3’ of the immunoglobulin heavy chain (IgH) genes (gift from Dr. Kathleen McGuire and Dr. Sanjeev Galande), as described previously^17–19^. HEK293T/17 cells were transfected with firefly luciferase reporter constructs and a Renilla luciferase (Rluc) normalization control (pGL4.74; Promega) in a ratio of 50:1, and with pYFP-SATB1 (WT or variant) or empty control vector (pYFP). After 48 h, firefly luciferase and Rluc activity was measured using the Dual-Luciferase Reporter Assay system (Promega) at the Infinite M Plex Microplate reader (Tecan).

### BRET saturation assays

BRET assays were performed as previously described^51^. HEK293T/17 cells were transfected in white clear-bottomed 96-well plates with increasing molar ratios of YFP-fusion proteins and constant amounts of Rluc-fusion proteins (donor/acceptor ratios of 1/0.5, 1/1, 1/2, 1/3, 1/6, 1/9). YFP and Rluc fused to a C-terminal nuclear localization signal were used as control proteins. After 48 h, medium was replaced with phenol red-free DMEM, supplemented with 10% fetal bovine serum (both Invitrogen), containing 60 μM EnduRen Live Cell Substrate (Promega). After incubation for 4h at 37°C, measurements were taken in live cells with an Infinite M200PRO Microplate reader (Tecan) using the Blue1 and Green1 filters. Corrected BRET ratios were calculated with the following formula: [Green1(_experimental condition_)/Blue1(_experimental condition_)] — [Green1(_control condition_)/Blue1(_control condition_)], with only the Rluc control protein expressed in the control condition. YFP fluorescence was measured separately (Ex: 505 nm, Em: 545 nm) to quantify expression of the YFP-fusion proteins. Curve fitting was done with a non-linear regression equation assuming a single binding site using GraphPad Prism Software, after plotting the corrected BRET ratios against the ratio of total luminescence / total YFP fluorescence.

### Immunoblotting and gel-shift assays

Whole-cell lysates were collected by treatment with lysis buffer 48 h post-transfection. For immunoblotting, cells were lysed in 1x RIPA buffer (Cell Signalling) with 1% PMSF and protease inhibitor cocktail (Roche). For gel-shift assays^54^, cells were lysed in 1x RIPA buffer with 1% PMSF, protease inhibitor cocktail and 50 μM ubiquitin/ubiquitin-like isopeptidases inhibitor PR-619 (Sigma). Samples were incubated for 20 min at 4°C followed by centrifugation for 30 min at 12,000 rpm at 4°C. Proteins were resolved on 4-15% Mini-PROTEAN TGX Precast Gels (Bio-Rad) and transferred onto polyvinylidene fluoride membranes using a TransBlot Turbo Blotting system (Bio-Rad). Membranes were blocked in 5% milk for 1 h at room temperature and then probed with mouse-anti-EGFP (for pYFP constructs; 1:8000; Clontech, 632380) or mouse-anti-V5 tag (1:2000; Genetex, GTX42525). Next, membranes were incubated with HRP-conjugated goat-anti-mouse IgG (1:2000; Bio-Rad) for 1 h at room temperature. Bands were visualized with Novex ECL Chemiluminescent Substrate Reagent (Invitrogen) using a ChemiDoc XRS + System (Bio-Rad). Equal protein loading was confirmed by probing with mouse-anti-β-actin antibody (1:10,000; Sigma, A5441).

### Fluorescence-based quantification of protein stability

Cells were transfected in triplicate in clear-bottomed black 96-well plates with YFP-tagged SATB1 variants. After 24 h, MG132 (R&D Systems) was added at a final concentration of 10 μM, and cycloheximide (Sigma) at 50 μg/ml. Cells were incubated at 37°C with 5% CO_2_ in the Infinite M200PRO microplate reader (Tecan), and the fluorescence intensity of YFP (Ex: 505 nm, Em: 545 nm) was measured over 24 h at 3 h intervals.

### Statistical analyses of cell-based functional assays

Statistical analyses for cell-based functional assays were done using a one-or two-way ANOVA followed by a Bonferroni *post-hoc* test, with GraphPad Prism Software. Statistical analyses for FRAP and BRET data were performed on values derived from fitted curves of individual recordings or independent experiments respectively.

## References

1. Wang, Z., Yang, X., Chu, X., Zhang, J., Zhou, H., Shen, Y., and Long, J. (2012). The structural basis for the oligomerization of the N-terminal domain of SATB1. Nucleic Acids Res 40, 4193–4202.

2. Alvarez, J.D., Yasui, D.H., Niida, H., Joh, T., Loh, D.Y., and Kohwi-Shigematsu, T. (2000). The MAR-binding protein SATB1 orchestrates temporal and spatial expression of multiple genes during T-cell development. Genes Dev 14, 521–535.

3. Cai, S., Lee, C.C., and Kohwi-Shigematsu, T. (2006). SATB1 packages densely looped, transcriptionally active chromatin for coordinated expression of cytokine genes. Nat Genet 38, 1278–1288.

4. Kitagawa, Y., Ohkura, N., Kidani, Y., Vandenbon, A., Hirota, K., Kawakami, R., Yasuda, K., Motooka, D., Nakamura, S., Kondo, M., et al. (2017). Guidance of regulatory T cell development by Satb1-dependent super-enhancer establishment. Nat Immunol 18, 173–183.

5. Satterstrom, F.K., Kosmicki, J.A., Wang, J., Breen, M.S., De Rubeis, S., An, J.Y., Peng, M., Collins, R., Grove, J., Klei, L., et al. (2020). Large-Scale Exome Sequencing Study Implicates Both Developmental and Functional Changes in the Neurobiology of Autism. Cell 180, 568–584.e523.

6. Kaplanis, J., Samocha, K.E., Wiel, L., Zhang, Z., Arvai, K.J., Eberhardt, R.Y., Gallone, G., Lelieveld, S.H., Martin, H.C., McRae, J.F., et al. (2020). Evidence for 28 genetic disorders discovered by combining healthcare and research data. Nature.

7. Sobreira, N., Schiettecatte, F., Valle, D., and Hamosh, A. (2015). GeneMatcher: a matching tool for connecting investigators with an interest in the same gene. Hum Mutat 36, 928–930.

8. Thompson, R., Johnston, L., Taruscio, D., Monaco, L., Beroud, C., Gut, I.G., Hansson, M.G., t Hoen, P.B., Patrinos, G.P., Dawkins, H., et al. (2014). RD-Connect: an integrated platform connecting databases, registries, biobanks and clinical bioinformatics for rare disease research. J Gen Intern Med 29 Suppl 3, S780–787.

9. Firth, H.V., Richards, S.M., Bevan, A.P., Clayton, S., Corpas, M., Rajan, D., Van Vooren, S., Moreau, Y., Pettett, R.M., and Carter, N.P. (2009). DECIPHER: Database of Chromosomal Imbalance and Phenotype in Humans Using Ensembl Resources. Am J Hum Genet 84, 524–533.

10. Karczewski, K.J., Francioli, L.C., Tiao, G., Cummings, B.B., Alföldi, J., Wang, Q., Collins, R.L., Laricchia, K.M., Ganna, A., Birnbaum, D.P., et al. (2019). Variation across 141,456 human exomes and genomes reveals the spectrum of loss-of-function intolerance across human protein-coding genes. bioRxiv, 531210.

11. Lelieveld, S.H., Reijnders, M.R., Pfundt, R., Yntema, H.G., Kamsteeg, E.J., de Vries, P., de Vries, B.B., Willemsen, M.H., Kleefstra, T., Lohner, K., et al. (2016). Meta-analysis of 2,104 trios provides support for 10 new genes for intellectual disability. Nat Neurosci 19, 1194–1196.

12. Lelieveld, S.H., Wiel, L., Venselaar, H., Pfundt, R., Vriend, G., Veltman, J.A., Brunner, H.G., Vissers, L., and Gilissen, C. (2017). Spatial Clustering of de Novo Missense Mutations Identifies Candidate Neurodevelopmental Disorder-Associated Genes. Am J Hum Genet 101, 478–484.

13. Kaufman, L., and Rousseeuw, P.J. (1987). Clustering by means of medoids. In. (

14. Köhler, S., Carmody, L., Vasilevsky, N., Jacobsen, J.O.B., Danis, D., Gourdine, J.P., Gargano, M., Harris, N.L., Matentzoglu, N., McMurry, J.A., et al. (2019). Expansion of the Human Phenotype Ontology (HPO) knowledge base and resources. Nucleic Acids Res 47, D1018–d1027.

15. Reijnders, M.R.F., Miller, K.A., Alvi, M., Goos, J.A.C., Lees, M.M., de Burca, A., Henderson, A., Kraus, A., Mikat, B., de Vries, B.B.A., et al. (2018). De Novo and Inherited Loss-of-Function Variants in TLK2: Clinical and Genotype-Phenotype Evaluation of a Distinct Neurodevelopmental Disorder. Am J Hum Genet 102, 1195–1203.

16. Walter, K., Min, J.L., Huang, J., Crooks, L., Memari, Y., McCarthy, S., Perry, J.R., Xu, C., Futema, M., Lawson, D., et al. (2015). The UK10K project identifies rare variants in health and disease. Nature 526, 82–90.

17. Pavan Kumar, P., Purbey, P.K., Sinha, C.K., Notani, D., Limaye, A., Jayani, R.S., and Galande, S. (2006). Phosphorylation of SATB1, a global gene regulator, acts as a molecular switch regulating its transcriptional activity in vivo. Mol Cell 22, 231–243.

18. Kumar, P.P., Purbey, P.K., Ravi, D.S., Mitra, D., and Galande, S. (2005). Displacement of SATB1-bound histone deacetylase 1 corepressor by the human immunodeficiency virus type 1 transactivator induces expression of interleukin-2 and its receptor in T cells. Mol Cell Biol 25, 1620–1633.

19. Siebenlist, U., Durand, D.B., Bressler, P., Holbrook, N.J., Norris, C.A., Kamoun, M., Kant, J.A., and Crabtree, G.R. (1986). Promoter region of interleukin-2 gene undergoes chromatin structure changes and confers inducibility on chloramphenicol acetyltransferase gene during activation of T cells. Mol Cell Biol 6, 3042–3049.

20. MacDermot, K.D., Bonora, E., Sykes, N., Coupe, A.M., Lai, C.S., Vernes, S.C., Vargha-Khadem, F., McKenzie, F., Smith, R.L., Monaco, A.P., et al. (2005). Identification of FOXP2 truncation as a novel cause of developmental speech and language deficits. Am J Hum Genet 76, 1074–1080.

21. Lindeboom, R.G.H., Vermeulen, M., Lehner, B., and Supek, F. (2019). The impact of nonsense-mediated mRNA decay on genetic disease, gene editing and cancer immunotherapy. Nat Genet.

22. Tan, J.A., Sun, Y., Song, J., Chen, Y., Krontiris, T.G., and Durrin, L.K. (2008). SUMO conjugation to the matrix attachment region-binding protein, special AT-rich sequence-binding protein-1 (SATB1), targets SATB1 to promyelocytic nuclear bodies where it undergoes caspase cleavage. J Biol Chem 283, 18124–18134.

23. Haijes, H.A., Koster, M.J.E., Rehmann, H., Li, D., Hakonarson, H., Cappuccio, G., Hancarova, M., Lehalle, D., Reardon, W., Schaefer, G.B., et al. (2019). De Novo Heterozygous POLR2A Variants Cause a Neurodevelopmental Syndrome with Profound Infantile-Onset Hypotonia. Am J Hum Genet 105, 283–301.

24. O’Donnell-Luria, A.H., Pais, L.S., Faundes, V., Wood, J.C., Sveden, A., Luria, V., Abou Jamra, R., Accogli, A., Amburgey, K., Anderlid, B.M., et al. (2019). Heterozygous Variants in KMT2E Cause a Spectrum of Neurodevelopmental Disorders and Epilepsy. Am J Hum Genet 104, 1210–1222.

25. Reynhout, S., Jansen, S., Haesen, D., van Belle, S., de Munnik, S.A., Bongers, E., Schieving, J.H., Marcelis, C., Amiel, J., Rio, M., et al. (2019). De Novo Mutations Affecting the Catalytic Calpha Subunit of PP2A, PPP2CA, Cause Syndromic Intellectual Disability Resembling Other PP2A-Related Neurodevelopmental Disorders. Am J Hum Genet 104, 139–156.

26. Reijnders, M.R.F., Ansor, N.M., Kousi, M., Yue, W.W., Tan, P.L., Clarkson, K., Clayton-Smith, J., Corning, K., Jones, J.R., Lam, W.W.K., et al. (2017). RAC1 Missense Mutations in Developmental Disorders with Diverse Phenotypes. Am J Hum Genet 101, 466–477.

27. Zarate, Y.A., Bosanko, K.A., Caffrey, A.R., Bernstein, J.A., Martin, D.M., Williams, M.S., Berry-Kravis, E.M., Mark, P.R., Manning, M.A., Bhambhani, V., et al. (2019). Mutation update for the SATB2 gene. Hum Mutat 40, 1013–1029.

28. Lee, J.S., Yoo, Y., Lim, B.C., Kim, K.J., Choi, M., and Chae, J.H. (2016). SATB2-associated syndrome presenting with Rett-like phenotypes. Clin Genet 89, 728–732.

29. Donnai, D., Tomlin, P.I., and Winter, R.M. (2005). Kohlschutter syndrome in siblings. Clin Dysmorphol 14, 123–126.

30. Retterer, K., Juusola, J., Cho, M.T., Vitazka, P., Millan, F., Gibellini, F., Vertino-Bell, A., Smaoui, N., Neidich, J., Monaghan, K.G., et al. (2016). Clinical application of whole-exome sequencing across clinical indications. Genet Med 18, 696–704.

31. de Ligt, J., Willemsen, M.H., van Bon, B.W., Kleefstra, T., Yntema, H.G., Kroes, T., Vulto-van Silfhout, A.T., Koolen, D.A., de Vries, P., Gilissen, C., et al. (2012). Diagnostic exome sequencing in persons with severe intellectual disability. N Engl J Med 367, 1921–1929.

32. DDD-study. (2015). Large-scale discovery of novel genetic causes of developmental disorders. Nature 519, 223–228.

33. Gueneau, L., Fish, R.J., Shamseldin, H.E., Voisin, N., Tran Mau-Them, F., Preiksaitiene, E., Monroe, G.R., Lai, A., Putoux, A., Allias, F., et al. (2018). KIAA1109 Variants Are Associated with a Severe Disorder of Brain Development and Arthrogryposis. Am J Hum Genet 102, 116–132.

34. Brunet, T., Radivojkov-Blagojevic, M., Lichtner, P., Kraus, V., Meitinger, T., and Wagner, M. (2020). Biallelic loss-of-function variants in RBL2 in siblings with a neurodevelopmental disorder. Ann Clin Transl Neurol 7, 390–396.

35. Yang, Y., Muzny, D.M., Xia, F., Niu, Z., Person, R., Ding, Y., Ward, P., Braxton, A., Wang, M., Buhay, C., et al. (2014). Molecular findings among patients referred for clinical whole-exome sequencing. Jama 312, 1870–1879.

36. Rentzsch, P., Witten, D., Cooper, G.M., Shendure, J., and Kircher, M. (2019). CADD: predicting the deleteriousness of variants throughout the human genome. Nucleic Acids Res 47, D886–d894.

37. Jaganathan, K., Kyriazopoulou Panagiotopoulou, S., McRae, J.F., Darbandi, S.F., Knowles, D., Li, Y.I., Kosmicki, J.A., Arbelaez, J., Cui, W., Schwartz, G.B., et al. (2019). Predicting Splicing from Primary Sequence with Deep Learning. Cell 176, 535–548.e524.

38. Wiel, L., Baakman, C., Gilissen, D., Veltman, J.A., Vriend, G., and Gilissen, C. (2019). MetaDome: Pathogenicity analysis of genetic variants through aggregation of homologous human protein domains. Hum Mutat 40, 1030–1038.

39. Boyd, A., Golding, J., Macleod, J., Lawlor, D.A., Fraser, A., Henderson, J., Molloy, L., Ness, A., Ring, S., and Davey Smith, G. (2013). Cohort Profile: the ‘children of the 90s’--the index offspring of the Avon Longitudinal Study of Parents and Children. Int J Epidemiol 42, 111–127.

40. Fraser, A., Macdonald-Wallis, C., Tilling, K., Boyd, A., Golding, J., Davey Smith, G., Henderson, J., Macleod, J., Molloy, L., Ness, A., et al. (2013). Cohort Profile: the Avon Longitudinal Study of Parents and Children: ALSPAC mothers cohort. Int J Epidemiol 42, 97–110.

41. Wang, K., Li, M., and Hakonarson, H. (2010). ANNOVAR: functional annotation of genetic variants from high-throughput sequencing data. Nucleic Acids Res 38, e164.

42. Wang, J.Z., Du, Z., Payattakool, R., Yu, P.S., and Chen, C.F. (2007). A new method to measure the semantic similarity of GO terms. Bioinformatics 23, 1274–1281.

43. Deng, Y., Gao, L., Wang, B., and Guo, X. (2015). HPOSim: an R package for phenotypic similarity measure and enrichment analysis based on the human phenotype ontology. PLoS One 10, e0115692.

44. Yamasaki, K., Akiba, T., Yamasaki, T., and Harata, K. (2007). Structural basis for recognition of the matrix attachment region of DNA by transcription factor SATB1. Nucleic Acids Res 35, 5073–5084.

45. Guex, N., and Peitsch, M.C. (1997). SWISS-MODEL and the Swiss-PdbViewer: an environment for comparative protein modeling. Electrophoresis 18, 2714–2723.

46. Inoue, K., Hayashi, F., Yokoyama, S., RIKEN Structural Genomics/Proteomics Initiative (RSGI). (2005). Solution structure of the second CUT domain of human SATB2. In. (

47. Izumi, K., Yoshida, M., Hayashi, F., Hatta, R., Yokoyama, S., RIKEN Structural Genomics/Proteomics Initiative (RSGI). (2004). Solution structure of the homeodomain of KIAA1034 protein.

48. Iyaguchi, D., Yao, M., Watanabe, N., Nishihira, J., and Tanaka, I. (2007). DNA recognition mechanism of the ONECUT homeodomain of transcription factor HNF-6. Structure 15, 75–83.

49. Estruch, S.B., Graham, S.A., Quevedo, M., Vino, A., Dekkers, D.H.W., Deriziotis, P., Sollis, E., Demmers, J., Poot, R.A., and Fisher, S.E. (2018). Proteomic analysis of FOXP proteins reveals interactions between cortical transcription factors associated with neurodevelopmental disorders. Hum Mol Genet 27, 1212–1227.

50. Estruch, S.B., Graham, S.A., Deriziotis, P., and Fisher, S.E. (2016). The language-related transcription factor FOXP2 is post-translationally modified with small ubiquitin-like modifiers. Sci Rep 6, 20911.

51. Deriziotis, P., Graham, S.A., Estruch, S.B., and Fisher, S.E. (2014). Investigating protein protein interactions in live cells using bioluminescence resonance energy transfer. J Vis Exp.

52. Koulouras, G., Panagopoulos, A., Rapsomaniki, M.A., Giakoumakis, N.N., Taraviras, S., and Lygerou, Z. (2018). EasyFRAP-web: a web-based tool for the analysis of fluorescence recovery after photobleaching data. Nucleic Acids Res 46, W467–w472.

53. Kohze, R., Dieteren, C.E.J., Koopman, W.J.H., Brock, R., and Schmidt, S. (2017). Frapbot: An open-source application for FRAP data. Cytometry A 91, 810–814.

54. Jakobs, A., Koehnke, J., Himstedt, F., Funk, M., Korn, B., Gaestel, M., and Niedenthal, R. (2007). Ubc9 fusion-directed SUMOylation (UFDS): a method to analyze function of protein SUMOylation. Nat Methods 4, 245–250.

