## Supplementary Figures and Tables for "Mutation-specific pathophysiological mechanisms define different neurodevelopmental disorders associated with SATB1 dysfunction"

34. Fédération Hospitalo-Universitaire Médecine Translationnelle et Anomalies du Développement (TRANSLAD), Centre Hospitalier Universitaire Dijon, Dijon, France.
35. Department of Rehabilitation and Development, Randall Children's Hospital at Legacy Emanuel Medical Center, Portland, Oregon, USA.
36. Division of Child Neurology and Inherited Metabolic Diseases, Centre for Paediatrics and Adolescent Medicine, University Hospital Heidelberg, Heidelberg, Germany.
37. Department of Neuropediatrics, Tokyo Metropolitan Neurological Hospital, Fuchu, Tokyo, Japan.
38. Division of Allergy and Immunology, Northwell Health, Great Neck, NY, USA.
39. Departments of Medicine and Pediatrics, Donald and Barbara Zucker School of Medicine at Hofstra/Northwell, Hempstead, NY, USA.
40. Princess Máxima Center for Pediatric Oncology, Utrecht, The Netherlands.
41. Pediatrics & Genetics, Alpharetta, USA.
42. Department of Human Genetics, Yokohama City University Graduate School of Medicine, Yokohama, Kanagawa, Japan.
43. Yorkshire Regional Genetics Service, Chapel Allerton Hospital, Leeds, UK.
44. Division of Medical Genetics & Metabolism, Children’s Hospital of the King’s Daughters, Norfolk, VA, USA.
45. Department of Pediatrics, Eastern Virginia Medical School, Norfolk, VA, USA.
46. West of Scotland Centre for Genomic Medicine, Queen Elizabeth University Hospital, Glasgow, UK.
47. Department of Pediatrics, Showa University School of Medicine, Shinagawa-ku, Tokyo, Japan.
48. Zuidwester, Middelharnis, The Netherlands.
49. Mendelics Genomic Analysis, Sao Paulo, SP Brazil.
50. University of Sao Paulo, School of Medicine, Sao Paulo, SP Brazil.
51. CHU Rennes, Univ Rennes, CNRS, IGDR, Service de Génétique Clinique, Centre de Référence Maladies Rares CLAD-Ouest, ERN ITHACA, Hôpital Sud, Rennes, France.
52. Department of Molecular and Human Genetics, Baylor College of Medicine, Houston, TX, 77030, USA.
53. Baylor Genetics, Houston, Texas, 77021, USA.
54. Department of Pediatrics, Division of Genetics and Genomic Medicine, Washington University School of Medicine, St. Louis, MO, USA.
55. GeneDx, 207 Perry Parkway Gaithersburg, Maryland, USA.
56. Division of Pediatric Neurology, Duke University Medical Center, Durham, North Carolina, USA.
57. Department of Genetics, Penang General Hospital, Jalan Residensi, Georgetown, Penang, Malaysia.
58. Clinical Genetics, Guy's Hospital, Great Maze Pond, London, UK.
59. Department of Biological and Medical Sciences, Headington Campus, Oxford Brookes University, UK.
60. Clinical Genetics, St Michael's Hospital Bristol, University Hospitals Bristol NHS Foundation Trust, Bristol, UK.
61. Sheffield Clinical Genetics Service, Sheffield Children's Hospital, Sheffield, UK.
62. Clinical Genomics Department, Ambry Genetics, Aliso Viejo, CA, USA.
63. Medigenome, Swiss Institute of Genomic Medicine, Geneva, Switzerland.
64. Department of Genetics and Cell Biology, Faculty of Health Medicine Life Sciences, Maastricht University Medical Center+, Maastricht University, Maastricht, The Netherlands.
65. Department of Pediatrics, Division of Medical Genetics, Duke University Medical Center, Durham, North Carolina, USA.
66. Center for Pediatric Genomic Medicine, Children's Mercy Hospital, Kansas City, MO, USA.
67. Department of Pathology and Laboratory Medicine, Children's Mercy Hospital, Kansas City, MO, USA.
68. Institute of Neurogenomics, Helmholtz Zentrum München, Munich, Germany.
69. Department of Genetics, Children's Hospital of Eastern Ontario, Ottawa, Ontario, Canada.
70. The University of Kansas School of Medicine Salina Campus, Salina, USA.
71. Oxford Centre for Genomic Medicine, Oxford University Hospitals NHS Foundation Trust, Oxford, UK.
72. These authors contributed equally to this work.

* To whom correspondence should be addressed:
Prof. Dr. S.E. Fisher


Dr. L.E.L.M. Vissers


**Contents:**


| | |
|---|---|
| - Supplementary Figure 1 | Pedigrees of (suspected) mosaic families with *SATB1* variants. |
| - Supplementary Figure 2 | Amino acid sequence alignments of the CUT1, CUT2 and Homeobox domain of SATB1. |
| - Supplementary Figure 3 | Heterozygous (partial) gene deletions of the *SATB1* gene. |
| - Supplementary Figure 4 | Clinical evaluation of individuals with SATB1 variants. |
| - Supplementary Figure 5 | Grouped HPO features based on semantic similarity and clustering results per individual. |
| - Supplementary Figure 6 | Overexpression of SATB1 missense variants as YFP-fusion proteins. |
| - Supplementary Figure 7 | MetaDome analysis of the *SATB1* missense variants. |
| - Supplementary Figure 8 | Functional characterization of the SATB1 p.R410* variant. |
| - Supplementary Figure 9 | Overexpression of SATB1 NMD-escaping PTVs as YFP-fusion proteins. |
| - Supplementary Figure 10 | SUMOylation of SATB1 protein truncating variants escaping NMD. |
| - Supplementary Figure 11 | The SATB2 p.E396Q missense variant has comparable effects on protein functions as the p.E407G and p.E530K/Q SATB1 variants affecting equivalent positions. |
| - Supplementary Figure 12 | Missense variants identified in individuals with NDD displayed in an amino acid sequence alignment of SATB2 and SATB1. |

| | |
|---|---|
| - Supplementary Table 2 | Splice-AI predictions for missense variants at intron-exon or exon-intron junctions. |
| - Supplementary Table 3 | Phenotypic information of individuals from the UK10K cohort with rare *SATB1* missense variants. |
| - Supplementary Table 4 | NMD efficacy predictions for SATB1 truncating variants. |
| - Supplementary Table 5 | Summary of clinical characteristics associated with *(de novo) SATB1* PTVs and (partial) gene deletions predicted to result in haploinsufficiency and PTVs in the last exon. |
| - Supplementary Table 7 | Primers for site-directed mutagenesis. |
| - Supplementary Table 8 | Primers for amplifying and subcloning human UBC9 (NM_194260.2) and SATB1 (NM_001131010.4). |
| - Supplementary Table 9 | Primers to amplify regions that include the SATB1 NMD escaping truncating variants used for testing for NMD. |

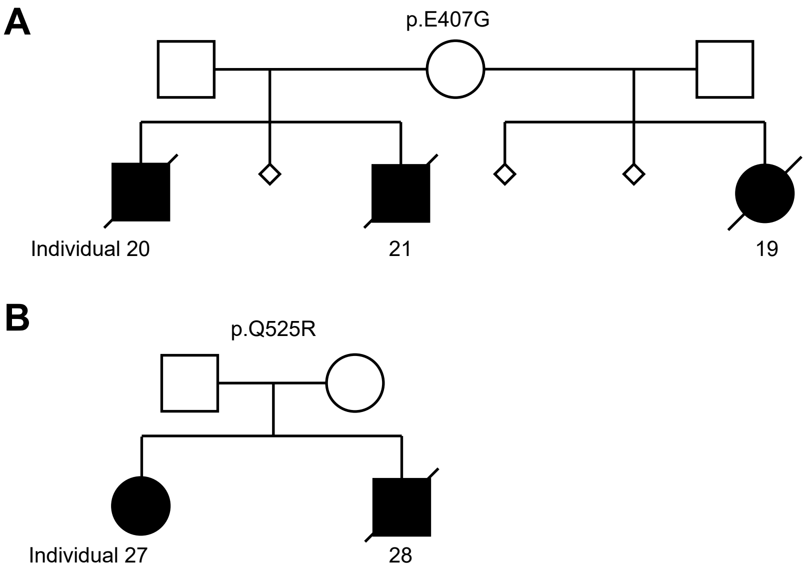


**Supplementary Figure 1. Pedigrees of (suspected) mosaic families with *SATB1* variants**. **A**) Pedigree of family with proband and siblings carrying a heterozygous SATB1 p.E407G variant. The mother presents the variant in 1 of 69 read in whole exome sequencing data, so the estimated percentage is 1.4 % in the peripheral blood. Karyotyping was normal. **B**) Pedigree of family with proband and sibling carrying a heterozygous SATB1 p.Q525R variant. Suspected mosaicism in one of the parents could not be confirmed with Sanger sequencing of DNA derived from peripheral blood. **A-B**) In both families, none of the pregnancies resulted in healthy offspring.

CUT1 domain

p.Q402R p.E407G/Q p.E413K p.Q420R

```
SP|Q01826|SATB1_HUMAN          LEQQVSTNTEVSSEIYQWVRDELKRAGISQAVFARVAFNRTQGLLSEILRKEEDPKTASQ 60
SP|Q60611|SATB1_MOUSE          LEQQVSTNTEVSSEIYQWVRDELKRAGISQAVFARVAFNRTQGLLSEILRKEEDPKTASQ 60
TR|Q5U2Y2|Q5U2Y2_RAT           LEQQVSTNTEVSSEIYQWVRDELKRAGISQAVFARVAFNRTQGLLSEILRKEEDPKTASQ 60
TR|A0A1D5PV61|A0A1D5PV61_CHICK LEQQVSTNTEVSSEIYQWVRDELKRAGISQAVFARVAFNRTQGLLSEILRKEEDPKTASQ 60
TR|F6W9B5|F6W9B5_XENTR         LEQQVSPNTEVSSDIYQWVRDELKRAGISQAVFARVAFNRTQGLLSEILRKEEDPKTASQ 60
SP|Q9UPW6|SATB2_HUMAN          KPEPTNSSVEVSPDIYQQVRDELKRASVSQAVFARVAFNRTQGLLSEILRKEEDPRTASQ 60
                                 : .. ..*** :*** ********.:***************************:****

SP|Q01826|SATB1_HUMAN          SLLVNLRAMQNFLQLPEAERDRIYQDER 88
SP|Q60611|SATB1_MOUSE          SLLVNLRAMQNFLQLPEAERDRIYQDER 88
TR|Q5U2Y2|Q5U2Y2_RAT           SLLVNLRAMQNFLQLPEAERDRIYQDER 88
TR|A0A1D5PV61|A0A1D5PV61_CHICK SLLVNLRAMQNFLQLPEAERDRIYQDER 88
TR|F6W9B5|F6W9B5_XENTR         SLLVNLRAMQNFLQLPEAERDRIYQDER 88
SP|Q9UPW6|SATB2_HUMAN          SLLVNLRAMQNFLNLPEVERDRIYQDER 88
                               *************:***.**********
```

CUT2 domain

p.Q525R p.E530K/Q/G

```
SP|Q01826|SATB1_HUMAN          NGKPENNTMNINASIYDEIQQEMKRAKVSQALFAKVAATKSQGWLCELLRWKEDPSPENR 60
SP|Q60611|SATB1_MOUSE          NGKPENNTMNINASIYDEIQQEMKRAKVSQALFAKVAATKSQGWLCELLRWKEDPSPENR 60
TR|Q5U2Y2|Q5U2Y2_RAT           NGKPENNTMNINASIYDEIQQEMKRAKVSQALFAKVAATKSQGWLCELLRWKEDPSPENR 60
TR|A0A1D5PV61|A0A1D5PV61_CHICK NGKTENNSMNINASIYDEIQQEMKRAKVSQALFAKVAATKSQGWLCELLRWKEDPSPENR 60
TR|F6W9B5|F6W9B5_XENTR         NGNLENCTMNINASIYDEIQQEMKRAKVSQALFAKVAASKSQGWLCELLRWKEDPSPENR 60
SP|Q9UPW6|SATB2_HUMAN          PIKVDGANINITAAIYDEIQQEMKRAKVSQALFAKVAANKSQGWLCELLRWKENPSPENR 60
                                 : :. .:**.*:************************.**************:******
```

p.E547K

```
SP|Q01826|SATB1_HUMAN          TLWENLSMIRRFLSLPQPERDAIYEQES 88
SP|Q60611|SATB1_MOUSE          TLWENLSMIRRFLSLPQPERDAIYEQES 88
TR|Q5U2Y2|Q5U2Y2_RAT           TLWENLSMIRRFLSLPQPERDAIYEQES 88
TR|A0A1D5PV61|A0A1D5PV61_CHICK TLWENLSMIRRFLSLPQPERDAIYEQES 88
TR|F6W9B5|F6W9B5_XENTR         TLWENLSMIRRFLSLPQTERDVIYEQES 88
SP|Q9UPW6|SATB2_HUMAN          TLWENLCTIRRFLNLPQHERDVIYEEES 88
                               ******. *****.*** ***.***:**
```

Homeobox domain

p.L682V

```
SP|Q01826|SATB1_HUMAN          ---TRPRTKISVEALGILQSFIQDVGLYPDEEAIQTLSAQLDLPKYTIIKFFQNQRYYLK 57
SP|Q60611|SATB1_MOUSE          ---TRPRTKISVEALGILQSFIQDVGLYPDEEAIQTLSAQLDLPKYTIIKFFQNQRYYLK 57
TR|Q5U2Y2|Q5U2Y2_RAT           RQKTRPRTKISVEALGILQSFIQDVGLYPDEEAIQTLSAQLDLPKYTIIKFFQNQRYYLK 60
TR|A0A1D5PV61|A0A1D5PV61_CHICK RQKPRPRTKISVEALGILQSFIQDVGLYPDEEAIQTLSAQLDLPKYTIIKFFQNQRYYLK 60
TR|F6W9B5|F6W9B5_XENTR         RQQPRPRTKISVEALGILQSFIQDVGLYPDEEAIQTLSAQLDLPKYTIIKFFQNQRYYLK 60
SP|Q9UPW6|SATB2_HUMAN          ---PRSRTKISLEALGILQSFIHDVGLYPDQEAIHTLSAQLDLPKHTIIKFFQNQRYHVK 57
                                   * *****:**********:*******:***:**********:***********::*

SP|Q01826|SATB1_HUMAN          HHG 60
SP|Q60611|SATB1_MOUSE          HHG 60
TR|Q5U2Y2|Q5U2Y2_RAT           HH- 62
TR|A0A1D5PV61|A0A1D5PV61_CHICK HH- 62
TR|F6W9B5|F6W9B5_XENTR         HH- 62
SP|Q9UPW6|SATB2_HUMAN          HHG 60
                               **
```

**Supplementary Figure 2. Amino acid sequence alignments of the CUT1, CUT2 and Homeobox domain of SATB1.** Amino acid sequences of the CUT1, CUT2 and Homeobox domain of human SATB1 (Q01826, UniProt) aligned to the mouse (Q60611), rat (Q5U2Y2), chicken (A0A1D5PV61) and Xenopus tropicalis (F6W9B5) sequences, and the sequences of the homolog domains in human SATB2 (Q9UPW6). Alignment was performed with Clustal Omega (1.2.4) with default settings using UnitProt alignment tool. Missense variants described in this study and identified in these functional domains are shaded in red.

1.05Mb deletion - chr3:17915162-18968823

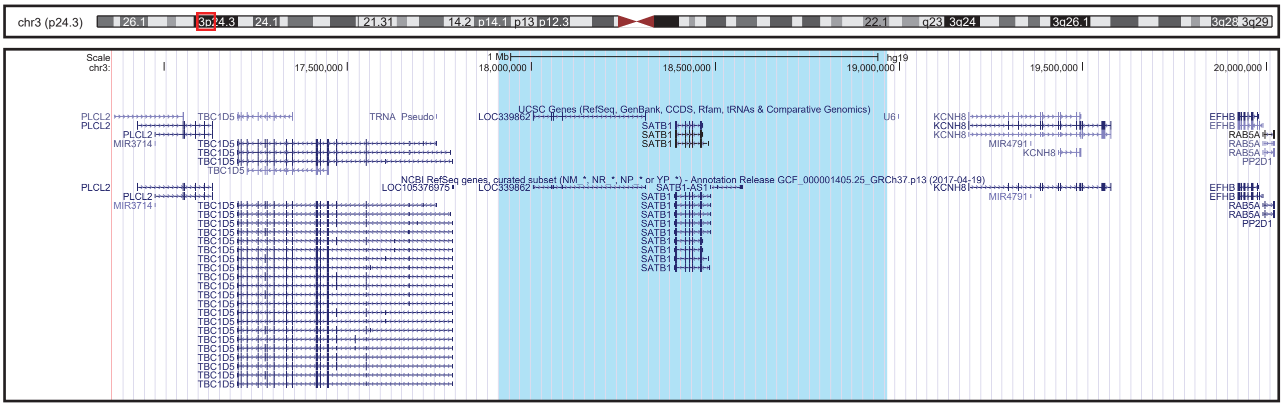


55Kb deletion - chr3:18,376,866-18,432,504

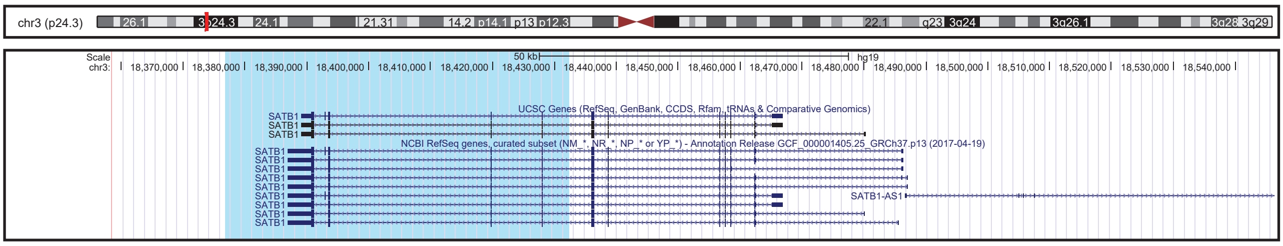


**Supplementary Figure 3. Heterozygous (partial) gene deletions of the *SATB1* gene.** Genome overviews of two reported heterozygous deletions that include the *SATB1* gene, generated in the UCSC Genome Browser (assembly Feb. 2009 GRCh37/hg19). The deleted regions are shaded in red in the chromosome ideogram, and in light blue in the genome overview.

**A-D**

Facial photographs not available in this preprint

**E**

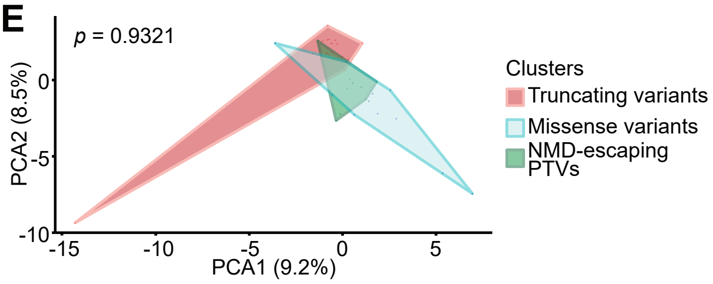


**Supplementary Figure 4. Clinical evaluation of individuals with *SATB1* variants.** A) Side view photographs, depicting prominent ears (individual 4, 8, 14, 17, 19, 34, 35), with thickened helices (individual 8, 14, 17, 19, 33, 34, 35), and retrognathia (individual 8, 14, 17, 19, 27, 34). B) Additional photograph of teeth. No evident enamel or dental positioning problems in individual 8 and 14, although missing molars (individual 8) and malformed teeth (individual 14) are reported. Lower teeth of individual 28: discoloration, malpositioning and teeth decay. C) Photographs of hands and feet. Features include contractures resulting from spasticity (individual 17), tapered fingers (individual 13, 14, 23, 35), short broad fingers (individual 13, 14, 23), clinodactyly of 5th finger (individual 9), overlapping 2nd toe (individual 35) or 4th toe (individual 9) and broad feet with short toes and small toe nails (individual 13, 14, 23). **D**) Computational average of facial photographs of 24 individuals with *SATB1* variants. **E**) Plot of Partitioning Around Medoids clustering analysis on clustered clinical data (HPO) showing no significant distinctions between individuals with missense variants, individuals with truncating variants and deletions, and individuals with NMD-escaping truncating variants

**A**

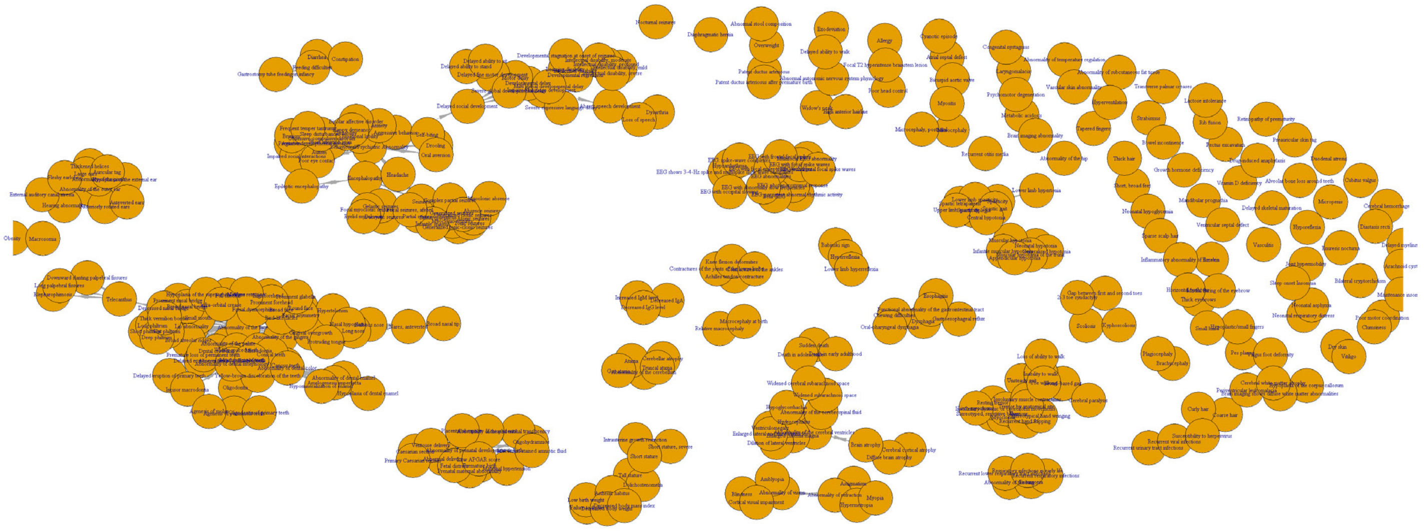

**B**

| Identifier | Variant type | 2_cluster _pred | 2_correct | 3_cluster _pred | 3_correct |
|---|---|---|---|---|---|
| Individual1 | PTV_non_last_exon | PTV | CORRECT | PTV_last_exon | INCORRECT |
| Individual2 | PTV_non_last_exon | PTV | CORRECT | PTV_non_last_exon | CORRECT |
| Individual3 | PTV_non_last_exon | PTV | CORRECT | PTV_last_exon | INCORRECT |
| Individual4 | PTV_non_last_exon | PTV | CORRECT | PTV_last_exon | INCORRECT |
| Individual5 | PTV_non_last_exon | Missense | INCORRECT | Missense | INCORRECT |
| Individual6 | PTV_non_last_exon | PTV | CORRECT | PTV_last_exon | INCORRECT |
| Individual7 | PTV_non_last_exon | Missense | INCORRECT | PTV_last_exon | INCORRECT |
| Individual8 | PTV_last_exon | PTV | CORRECT | PTV_last_exon | CORRECT |
| Individual9 | PTV_last_exon | PTV | CORRECT | PTV_last_exon | CORRECT |
| Individual10 | PTV_last_exon | PTV | CORRECT | PTV_last_exon | CORRECT |
| Individual11 | PTV_last_exon | PTV | CORRECT | PTV_non_last_exon | INCORRECT |
| Individual12 | PTV_last_exon | PTV | CORRECT | PTV_last_exon | CORRECT |

| Identifier | Variant type | 2_cluster _pred | 2_correct | 3_cluster _pred | 3_correct |
|---|---|---|---|---|---|
| Individual13 | Missense | PTV | INCORRECT | PTV_last_exon | INCORRECT |
| Individual14 | Missense | PTV | INCORRECT | PTV_last_exon | INCORRECT |
| Individual15 | Missense | PTV | INCORRECT | PTV_last_exon | INCORRECT |
| Individual17 | Missense | Missense | CORRECT | Missense | CORRECT |
| Individual18 | Missense | PTV | INCORRECT | PTV_last_exon | INCORRECT |
| Individual19 | Missense | Missense | CORRECT | Missense | CORRECT |
| Individual20 | Missense | Missense | CORRECT | Missense | CORRECT |
| Individual21 | Missense | Missense | CORRECT | Missense | CORRECT |
| Individual23 | Missense | PTV | INCORRECT | PTV_last_exon | INCORRECT |
| Individual24 | Missense | Missense | CORRECT | Missense | CORRECT |
| Individual25 | Missense | Missense | CORRECT | PTV_non_last_exon | INCORRECT |
| Individual26 | Missense | Missense | CORRECT | Missense | CORRECT |
| Individual27 | Missense | Missense | CORRECT | Missense | CORRECT |
| Individual28 | Missense | Missense | CORRECT | Missense | CORRECT |
| Individual29 | Missense | Missense | CORRECT | Missense | CORRECT |
| Individual30 | Missense | Missense | CORRECT | Missense | CORRECT |
| Individual31 | Missense | Missense | CORRECT | PTV_non_last_exon | INCORRECT |
| Individual33 | Missense | Missense | CORRECT | PTV_non_last_exon | INCORRECT |
| Individual34 | Missense | Missense | CORRECT | PTV_non_last_exon | INCORRECT |
| Individual35 | Missense | PTV | INCORRECT | PTV_last_exon | INCORRECT |
| Individual36 | Missense | PTV | INCORRECT | PTV_last_exon | INCORRECT |
| Individual37 | Missense | Missense | CORRECT | Missense | CORRECT |
| Individual38 | Missense | Missense | CORRECT | Missense | CORRECT |
| Individual39 | Missense | Missense | CORRECT | PTV_non_last_exon | INCORRECT |
| Individual40 | Missense | PTV | INCORRECT | PTV_last_exon | INCORRECT |
| Individual42 | Missense | PTV | INCORRECT | PTV_last_exon | INCORRECT |
| **Correctly predicted individuals:** | | | **27** | | **17** |

**Supplementary Figure 5. Grouped HPO features based on semantic similarity and clustering results per individual. A**) The semantic similarity between all the HPO terms used in this cohort (356 features) was calculated using the Wang algorithm in the HPOsim package in R. HPO terms with at least a 0.5 similarity score were grouped and a new feature was created as a replacement, which was the sum of the grouped features. **B**) Individual HPO-based phenotypic clustering results for both analyses with two and three clusters.

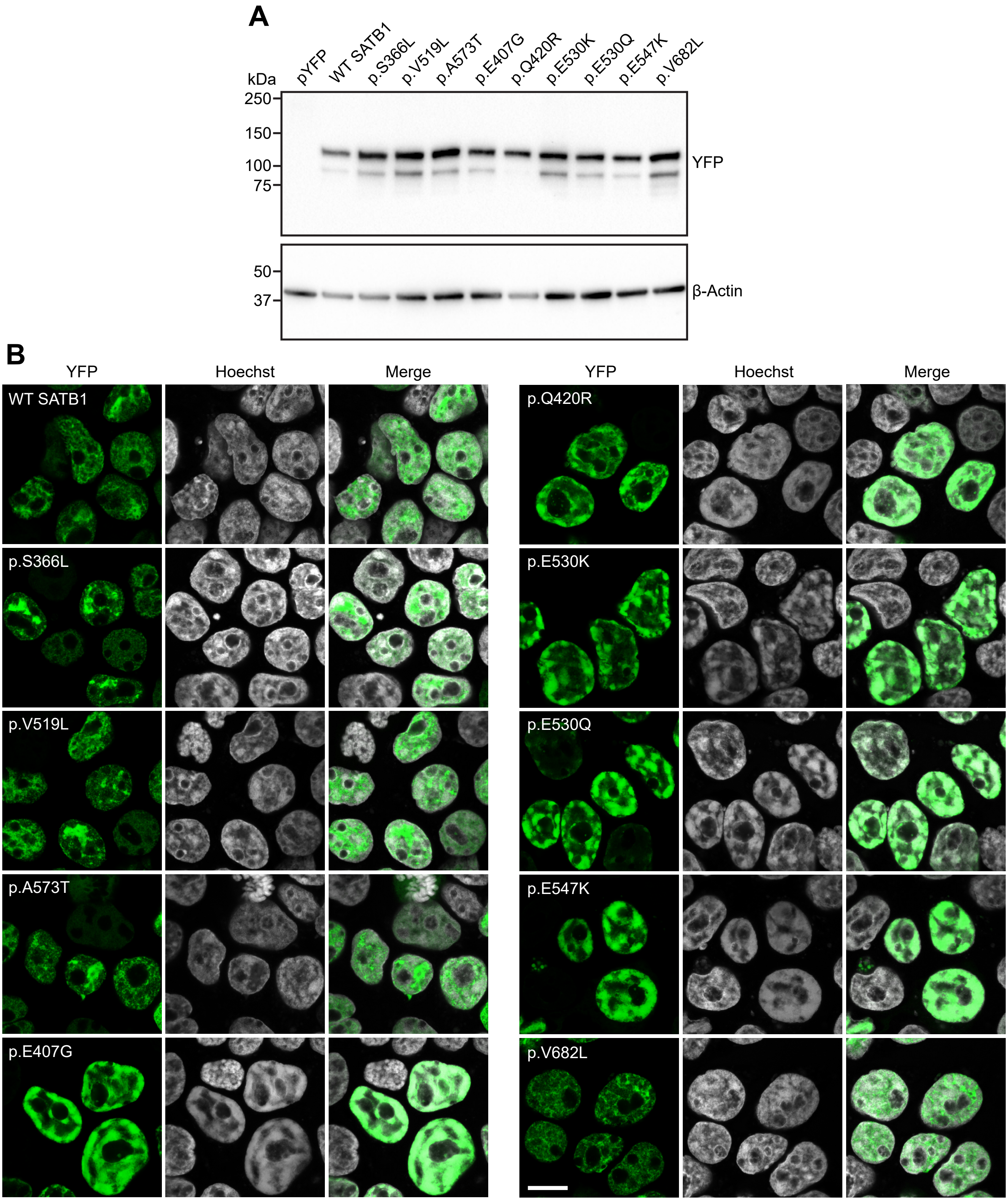


**Supplementary Figure 6. Overexpression of SATB1 missense variants as YFP-fusion proteins. A**) Immunoblot of whole-cell lysates expressing YFP-tagged SATB1 variants probed with anti-EGFP antibody. Expected molecular weight for all variants is ~115 kDa. The blot was probed for β-actin to ensure equal protein loading. **B**) Direct fluorescence micrographs of HEK293T/17 cells expressing YFP-SATB1 fusion proteins (green). Nuclei were stained with Hoechst 33342 (white). Scale bar = 10 µm.

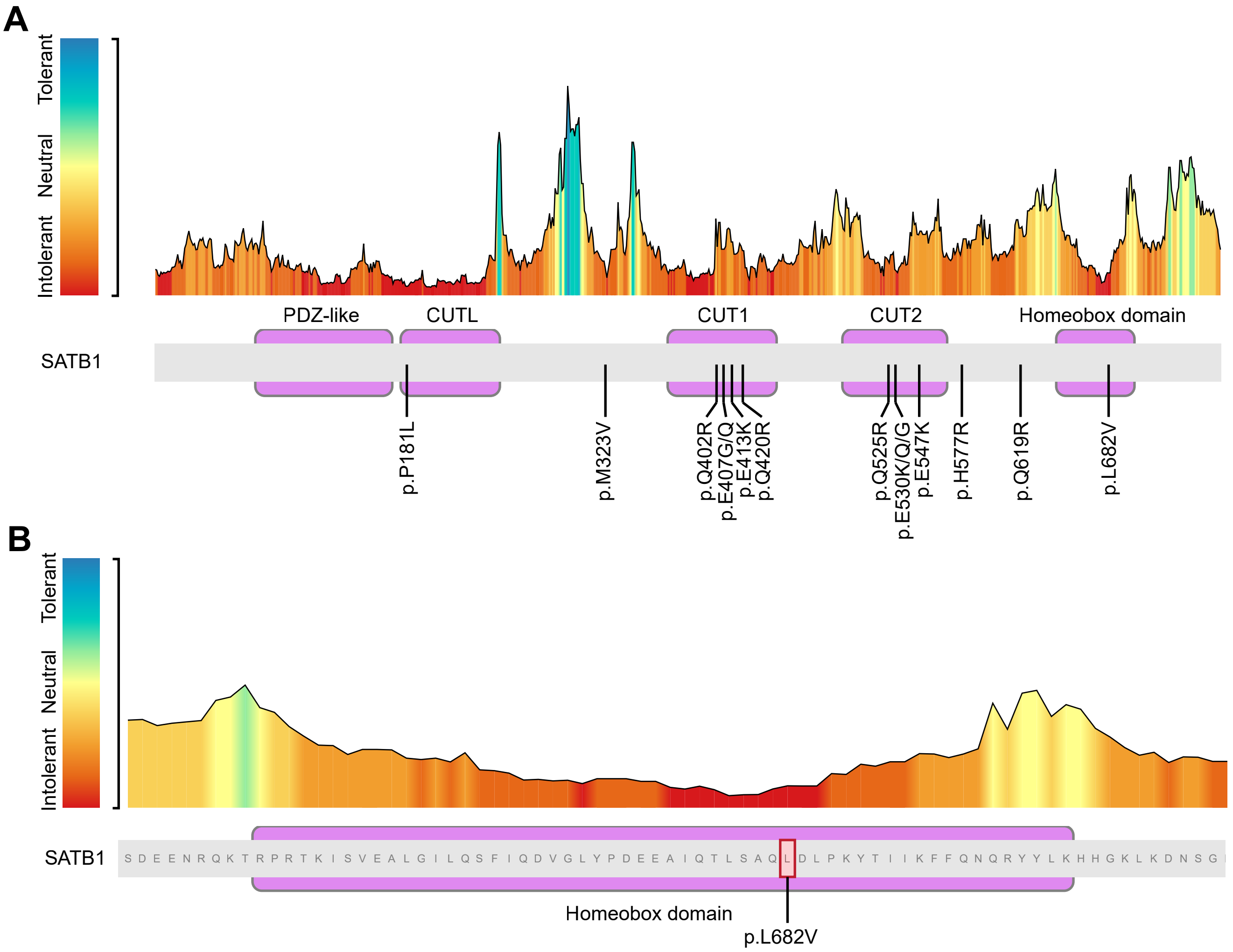


**C**

| Gene | Position | Variant | Residue change | Type | gnomAD Allele Frequency |
|---|---|---|---|---|---|
| HMX3 | chr10:124896963 | C>G | Leu>Val | missense | 0.000004 |
| POU2F2 | chr19:42599569 | G>C | Leu>Val | missense | 0.000004 |
| HOXC11 | chr12:54369087 | C>G | Leu>Val | missense | 0.000004 |
| HOXC10 | chr12:54383114 | A>G | Ile>Val | missense | 0.000004 |
| ZFHX4 | chr8:77767083 | C>A | Val>Val | synonymous | 0.000096 |
| POU6F1 | chr12:51584125 | G>C | Leu>Val | missense | 0.000004 |
| NKX1-2 | chr10:126136333 | G>C | Leu>Val | missense | 0.000009 |
| NOBOX | chr7:144097323 | C>T | Val>Val | synonymous | 0.000004 |
| ZFHX4 | chr8:77767083 | C>T | Val>Val | synonymous | 0.000008 |
| OTP | chr5:76932672 | T>C | Ile>Val | missense | 0.000070 |
| ISX | chr22:35478636 | A>G | Ile>Val | missense | 0.000004 |
| ZFHX3 | chr16:72828547 | C>T | Val>Val | synonymous | 0.000004 |
| PAX3 | chr2:223096822 | G>A | Ala>Val | missense | 0.000004 |
| NANOGNB | chr12:7922891 | T>G | Phe>Val | missense | 0.000023 |
| NKX2-2 | chr20:21492890 | T>C | Ile>Val | missense | 0.000004 |

**Supplementary Figure 7. MetaDome analysis of the *SATB1* missense variants. A**) Overview of the SATB1 protein (transcript NM_001131010.2) tolerance landscape. All missense variants identified in affected individuals are indicated. **B**) Detailed overview of the SATB1 homeobox domain tolerance landscape, with the p.L682V variant indicated. **C**) Table listing all residue changes at positions equivalent to the SATB1 p.L682 position in homolog homeobox domain proteins that change to a valine. The GnomAD allele frequency is indicated.

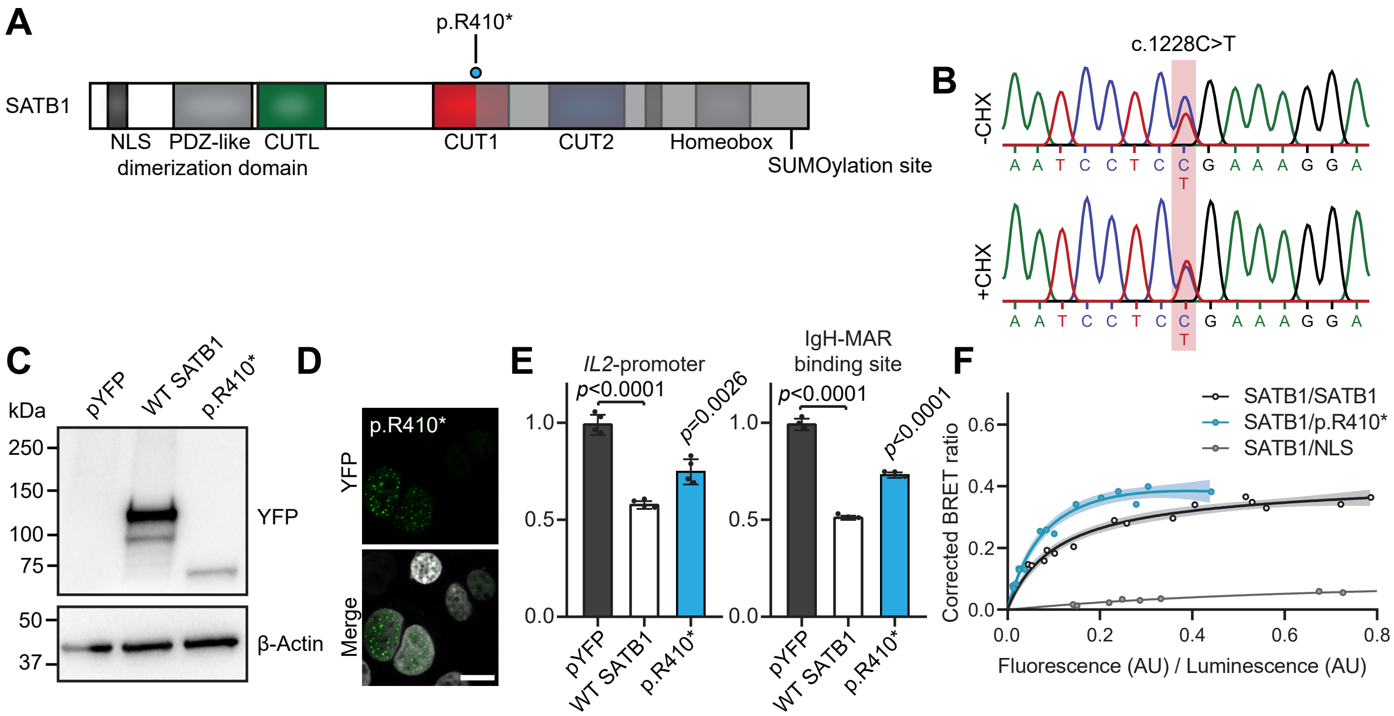


**Supplementary Figure 8. Functional characterization of the SATB1 p.R410* variant. A**) Schematic representation of SATB1 with the p.R410* variant labeled in cyan. **B**) Sanger sequencing traces of patient-derived EBV transformed lymphoblastoid cell lines treated with or without cycloheximide (CHX) to test for NMD. The mutated nucleotides are shaded in red. **C**) Immunoblot of whole-cell lysates expressing YFP-tagged SATB1 and p.R410* probed with anti-EGFP antibody. Expected molecular weight is SATB1: ~115 kDa, p.R410*: ~75kDa. The blot was probed for β-actin to ensure equal protein loading. **D**) Direct fluorescence micrographs of HEK293T/17 cells expressing YFP-SATB1 p.R410* fusion proteins (green). Nuclei were stained with Hoechst 33342 (white). Scale bar = 10 µm. **E**) Luciferase reporter assays using reporter constructs containing the IL2 promoter region and the IgH matrix associated region (MAR) binding site. Values are expressed relative to the control (pYFP; black) and represent the mean ± S.E.M. ($n$ = 4 for IL2-promoter, $n$ = 3 for IgH-MAR binding site, $p$-values compared to wildtype (WT) SATB1 (white), one-way ANOVA and *post-hoc* Bonferroni test). **F**) BRET assays for SATB1 dimerization in live cells. The plot shows the mean BRET saturation curves ± 95% C.I. fitted using a non-linear regression equation assuming a single binding site ($y$ = BRETmax * $x$ / (BRET50 / $x$); GraphPad). The corrected BRET ratio is plotted against the ratio of fluorescence/luminescence (AU) to correct for expression level differences between conditions ($n$ = 3).

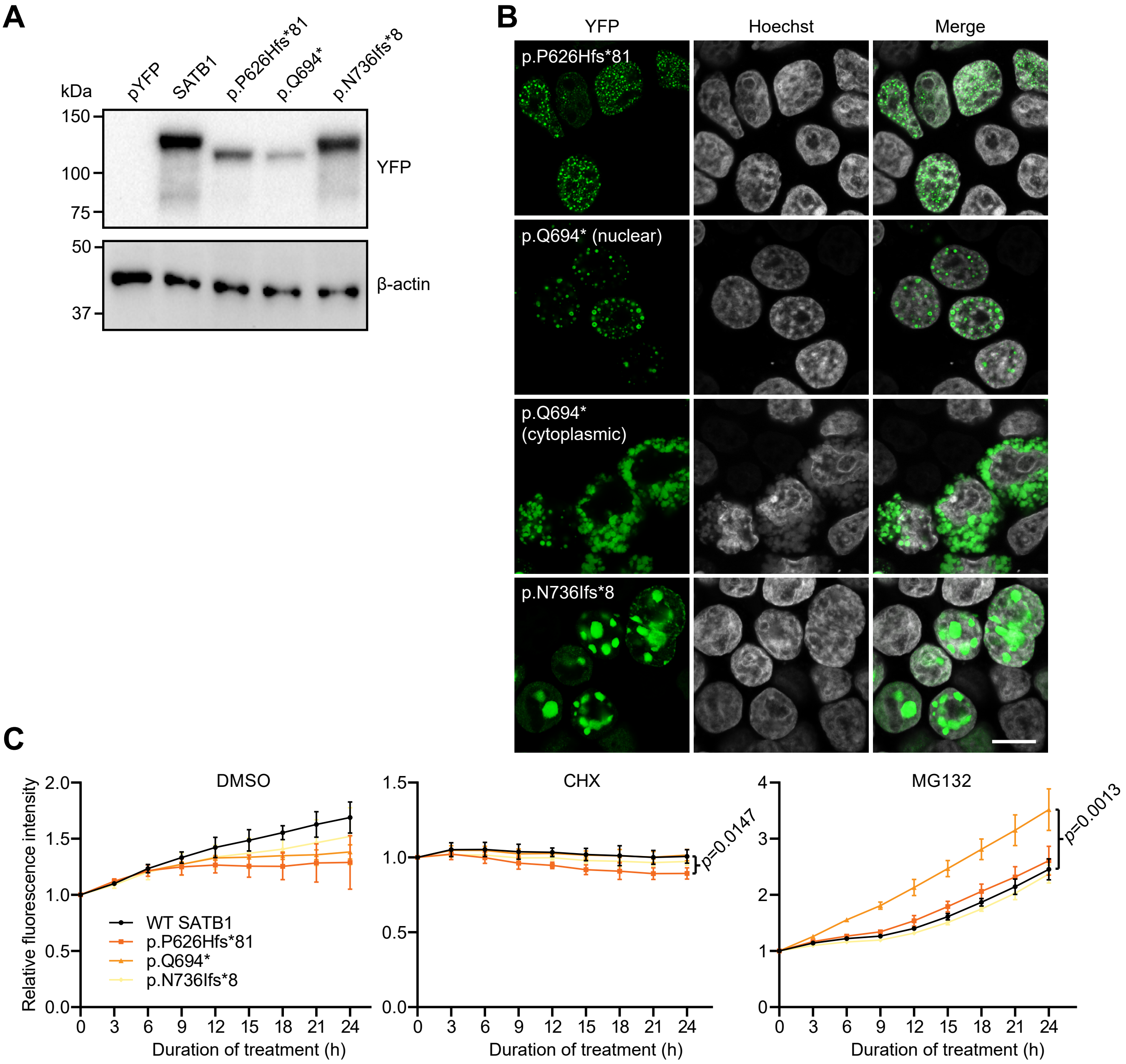


**Supplementary Figure 9. Overexpression of SATB1 NMD-escaping PTVs as YFP-fusion proteins. A**) Immunoblot of whole-cell lysates expressing YFP-tagged SATB1 variants probed with anti-EGFP antibody. Expected molecular weight: WT SATB1 = ~115 kDa, p.P626Hfs*81 = ~109 kDa, p.Q694* = ~107 kDa, p.N736Ifs*8 = ~113 kDa. The blot was probed for β-actin to ensure equal protein loading. **B**) Direct fluorescence imaging of HEK293T/17 cells expressing YFP-SATB1 fusion proteins (green). Nuclei were stained with Hoechst 33342 (white). Scale bar = 10 μm. **C**) Results of assay for protein stability of SATB1 NMD-escaping PTVs, using cycloheximide (CHX) to arrest protein synthesis, and MG132 to block protein degradation by the 26S proteasome complex. Values represent the mean protein expression levels of YFP-tagged SATB1 variants ± S.E.M. in live cells as measured by YFP fluorescence and expressed relative to the 0 h time point ($n$ = 3, two-way ANOVA for repeated measures with Geisser-Greenhouse correction, followed by a *post-hoc* Bonferroni test). Although p.P626Hfs*81 showed a slight but significant decrease in relative expression level after treatment with CHX, and p.Q694* showed a significant increase in relative expression level after treatment with MG132 when compared to WT SATB1, none of the variants tested showed both a decrease in levels after CHX treatment and an increase after MG132 treatment, which would be indicative of reduced protein stability.

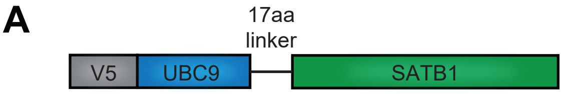


B

| Position K | Sequence | Best PS |
|---|---|---|
| K24 | SEMSNNVSDPKGPPAKIARLE | None |
| K29 | NVSDPKGPPAKIARLEQNGSP | None |
| K92 | HYENAIEYDCKEEHAEFVLVR | None |
| K175 | TLKIQLHSCPKLEDLPPEQWS | High |
| K198 | TVRNALKDLLKDMNQSSLAKE | None |
| K244 | QEFGRWYKHFKKTKDMMVEMD | None |
| K245 | EFGRWYKHFKKTKDMMVEMDS | None |
| K384 | EIYQWVRDELKRAGISQAVFA | None |
| K475 | STPPSRPPQVKTATIATERNG | None |
| K486 | TATIATERNGKPENNTMNINA | None |
| K523 | QALFAKVAATKSQGWLCELLR | None |
| K535 | QGWLCELLRWKEDPSPENRTL | None |
| K650 | ENRQKTRPRTKISVEALGILQ | None |
| K720 | SGLEVDVAEYKEEELLKDLEE | None |
| K726 | VAEYKEEELLKDLEESVQDKN | None |
| K744 | DKNTNTLFSVKLEEELSVEGN | High |

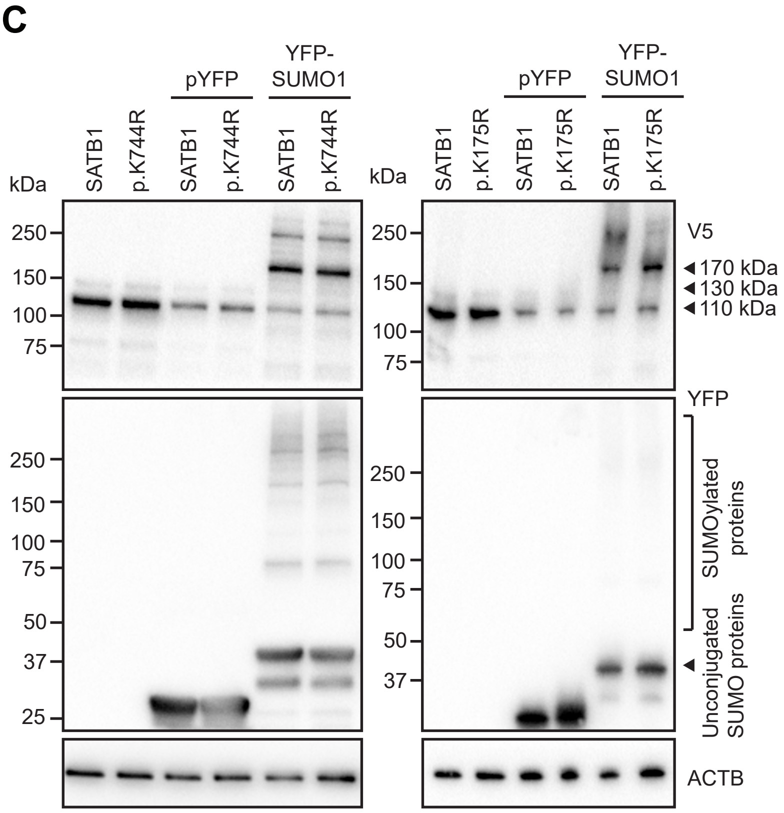


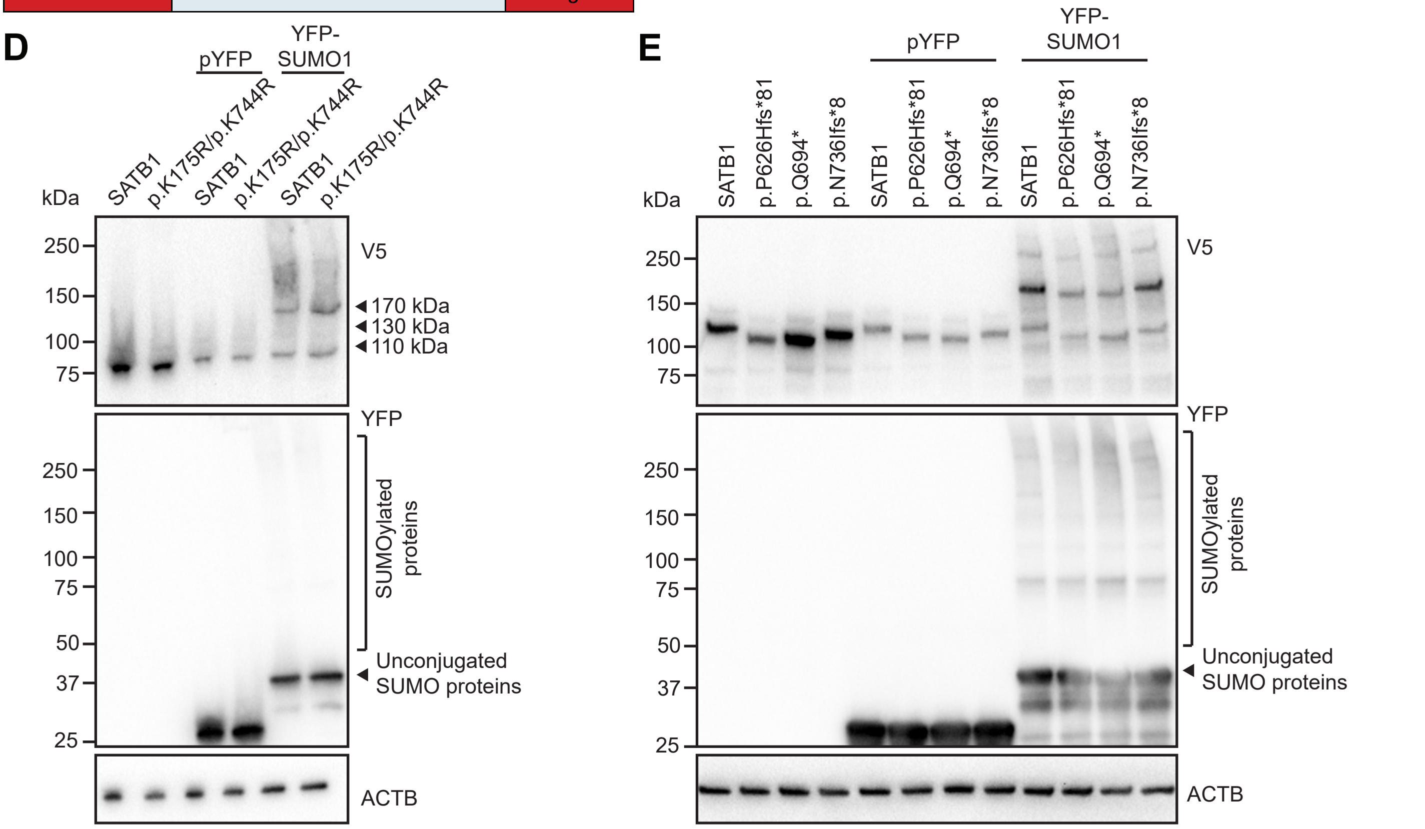


**Supplemenatry Figure 10. SUMOylation of SATB1 protein truncating variants escaping NMD. A**) Schematic representation of the UBC9-SATB1 fusion protein with an N-terminal V5 epitope tag. **B**) Prediction of putative SATB1 (Uniprot Q01826) SUMOylation sites using Joined Advanced SUMOylation Site and SIM Analyser (JASSA, www.jassa.fr/). JASSA uses a scoring system based on a Position Frequency Matrix derived from the alignment of experimental SUMOylation sites. K175 corresponds to a direct consensus site ([Ψ]-[K]-[x]-[α], with Ψ = A,F,I,L,M,P,V or W; α = D or E) with a high prediction score (PS), and K744 to a negatively charged amino acid-dependent SUMOylation site (NDSM, [Ψ]-[K]-[x]-[α]-[x]-$[α]_6$ with Ψ = A,F,I,L,M,P,V or W; 2 out of 6 α must be D or E) with a high PS. **C**) Gel shift assay for SATB1 SUMOylation. UBC9-SATB1 and a p.K175R or p.K744R mutant were expressed in HEK293T/17 cells together with a YFP-fusion of SUMO1. Top panel: western blot probed with anti-V5 antibody to detect UBC9-SATB1. The 110 kDa species is unmodified UBC9-SATB1. The 130 kDa species is UBC9-SATB1 modified with endogenous SUMO1. The 170 kDa species is UBC9-SATB1 modified with YFP-SUMO1. Middle panel: western blot probed with anti-YFP antibody, with unconjugated YFP-SUMO1 indicated with an arrow head. Higher molecular weight species are cellular proteins modified with YFP-SUMO1. Bottom panel: western blot probed with anti-β-actin to confirm equal protein loading. **D**) Gel-shift assay for SUMOylation of a SATB1 p.K175R/p.K744R double-mutant. **E**) Gel-shift assay for SUMOylation of SATB1 NMD escaping protein truncating variants.

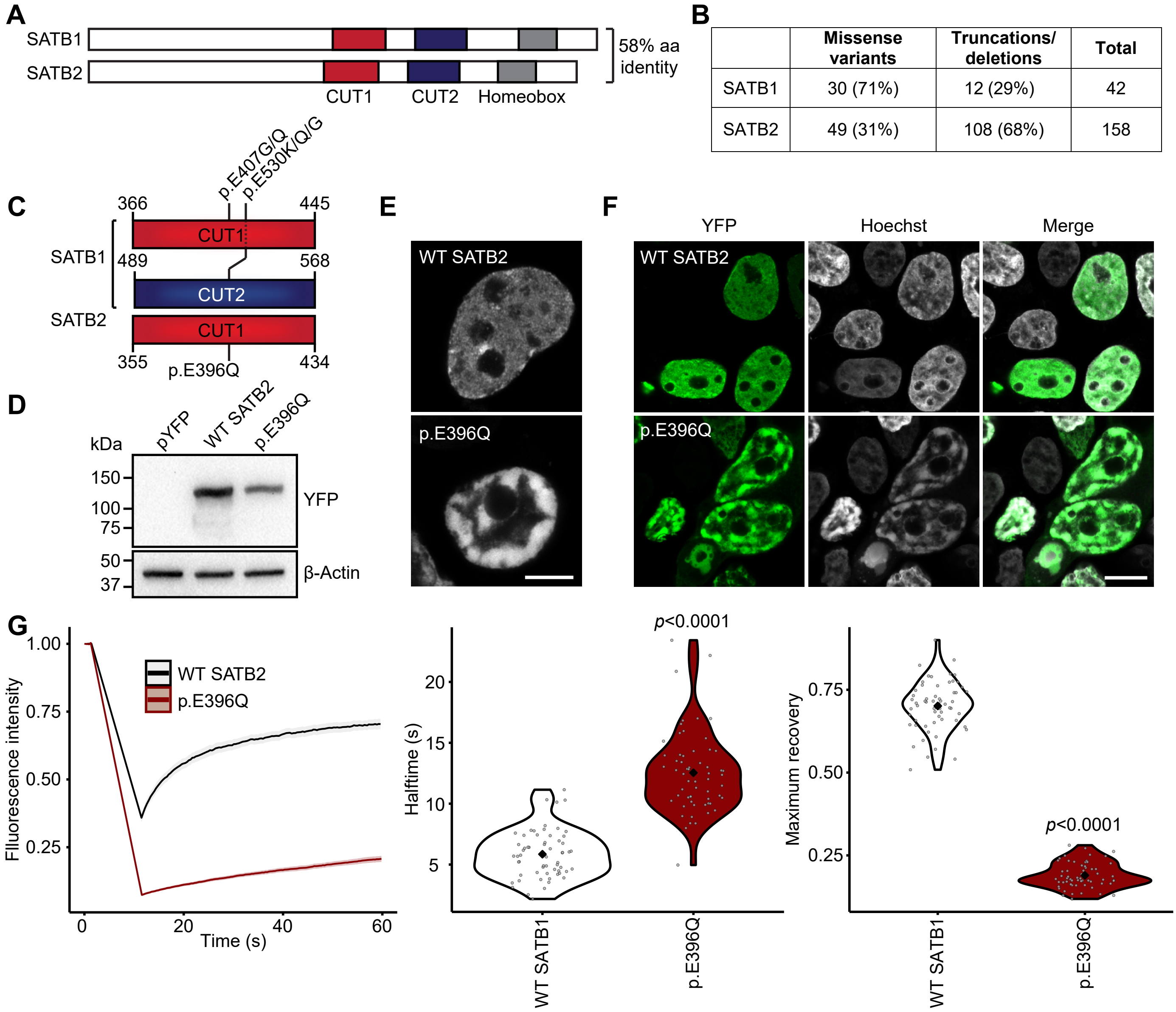


| | Missense variants | Truncations/ deletions | Total |
|---|---|---|---|
| SATB1 | 30 (71%) | 12 (29%) | 42 |
| SATB2 | 49 (31%) | 108 (68%) | 158 |

**Supplementary Figure 11. The SATB2 p.E396Q missense variant has comparable effects on protein functions as the p.E407G and p.E530K/Q SATB1 variants affecting equivalent positions. A**) SATB1 and SATB2 are highly conserved paralogs. **B**) In SATB1 more missense variants (71%) than truncations/deletions (29%) are observed, while for SATB2 the reverse is reported (31% versus 68% respectively). **C**) Schematic representation of SATB1 and SATB2 CUT DNA binding domains, with variants on equivalent positions indicated. **D**) Immunoblot of whole-cell lysates expressing YFP-tagged SATB2 and p.E396Q probed with anti-EGFP antibody. Expected molecular weight is ~112 kDa. The blot was probed for β-actin to ensure equal protein loading. **E**) Direct fluorescence super-resolution imaging of nuclei of HEK293T/17 cells expressing YFP-SATB2 fusion proteins. Scale bar = 5µm. **F**) Direct fluorescence imaging of HEK293T/17 cells expressing YFP-SATB2 fusion proteins (green). Nuclei were stained with Hoechst 33342 (white). Scale bar=10µm. **G)** FRAP experiments to assess the dynamics of SATB2 chromatin binding in live cells. Left, mean recovery curves ± 95% C.I. recorded in HEK293T/17 cells expressing YFP-SATB2 fusion proteins. Right, violin plots with median of the halftime and maximum recovery values based on single-term exponential curve fitting of individual recordings (*n* = 60 nuclei from three independent experiments, *p*-values compared to WT SATB2, unpaired t-test).

```
SP|Q9UPW6|SATB2_HUMAN MERRSESPCLRDSPDRRSGSPDVKGPPPVKVARLEQNGSPMGARGRPN------GA---- 50
SP|Q01826|SATB1_HUMAN MDHLNEATQGKEHSEMSNNVSDP-KGPPAKIARLEQNGSPLGRGRLGSTGAKMQGVPLKH 59
                      *:: .*:   ::  :  ..  *     **.*:*********:*      .       *.

SP|Q9UPW6|SATB2_HUMAN -----VAKAVGGLMIPVFCVVEQLDGSLEYDNREEHAEFVLVRKDVLFSQLVETALLALG 105
SP|Q01826|SATB1_HUMAN SGHLMKTNLRKGTMLPVFCVVEHYENAIEYDCKEEHAEFVLVRKDMLFNQLIEMALLSLG 119
                            ::   * *:*******: :.::*** :************:**.**:* ***:**

SP|Q9UPW6|SATB2_HUMAN YSHSSAAQAQGIIKLGRWNPLPLSYVTDAPDATVADMLQDVYHVVTLKIQLQSCSKLEDL 165
SP|Q01826|SATB1_HUMAN YSHSSAAQAKGLIQVGKWNPVPLSYVTDAPDATVADMLQDVYHVVTLKIQLHSCPKLEDL 179
                      *********:*:*::*:***:*****************************:** *****

SP|Q9UPW6|SATB2_HUMAN PAEQWNHATVRNALKELLKEMNQSTLAKECPLSQSMISSIVNSTYYANVSATKCQEFGRW 225
SP|Q01826|SATB1_HUMAN PPEQWSHTTVRNALKDLLKDMNQSSLAKECPLSQSMISSIVNSTYYANVSAAKCQEFGRW 239
                      * ***.*:*******:***:****:*************************:********

SP|Q9UPW6|SATB2_HUMAN YKKYKKIKVERVERENLSDYCVLGQRPMHLPNMNQLASLGKTNEQSPHSQIHHSTPIRNQ 285
SP|Q01826|SATB1_HUMAN YKHFKKTKDMMVEMDSLSELSQQGANHVN---FGQQPVPGNTAEQPPSPA-QLSHGSQPS 295
                      **::** *   ** :.**: .  * . ::   :.*    *:* ** *     : *   : .

SP|Q9UPW6|SATB2_HUMAN VPALQPIMSPGLLSPQLSPQLVRQQIAMAHLINQQIAVSRLLAHQHPQAINQQFLNHPPI 345
SP|Q01826|SATB1_HUMAN VRTPLPNLHPGLVSTPISPQLVNQQLVMAQLLNQQYAVNRLLAQQ---SLNQQYLNHPPP 352
                      * :  * : ***:*  :*****.**:.**:*:*** **.****:*    ::***:*****

SP|Q9UPW6|SATB2_HUMAN PRAVKPEP----TNSSVEVSPDIYQQVRDELKRASVSQAVFARVAFNRTQGLLSEILRKE 401
SP|Q01826|SATB1_HUMAN VSRSMNKPLEQQVSTNTEVSSEIYQWVRDELKRAGISQAVFARVAFNRTQGLLSEILRKE 412
                            :*    ..:..*** :*** ********.:************************

SP|Q9UPW6|SATB2_HUMAN EDPRTASQSLLVNLRAMQNFLNLPEVERDRIYQDERERSMNPNVSMVSSASSSPSSSRTP 461
SP|Q01826|SATB1_HUMAN EDPKTASQSLLVNLRAMQNFLQLPEAERDRIYQDERERSLNAASAMGPAPLISTPPSRPP 472
                      ***:*****************:***.*************:*   :*  :    *    ** *

SP|Q9UPW6|SATB2_HUMAN QAKTSTPTTDLPIKVDGANINITAAIYDEIQQEMKRAKVSQALFAKVAANKSQGWLCELL 521
SP|Q01826|SATB1_HUMAN QVKTATIATERNGKPENNTMNINASIYDEIQQEMKRAKVSQALFAKVAATKSQGWLCELL 532
                      *.**:* :*:   * :. .:**.*:***********************.**********

SP|Q9UPW6|SATB2_HUMAN RWKENPSPENRTLWENLCTIRRFLNLPQHERDVIYEEESR--HHHSERMQHVVQLPPEPV 579
SP|Q01826|SATB1_HUMAN RWKEDPSPENRTLWENLSMIRRFLSLPQPERDAIYEQESNAVHHHGDRPPHIIHVPAEQI 592
                      ****:************. *****.*** ***.***:**.   ***.:*  *::::* * :

SP|Q9UPW6|SATB2_HUMAN QVLHRQQSQPAKESS-----------------PPREEAPPPPPPTEDSCAKKPRSRTKIS 622
SP|Q01826|SATB1_HUMAN QQQQQQQQQQQQQQQQAPPPPQPQQQPQTGPRLPPRQPTVASPAESDEENRQKTRPRTKIS 652
                      *  ::**.*  ::..                 ***: :   *  :::.  :* * *****

SP|Q9UPW6|SATB2_HUMAN LEALGILQSFIHDVGLYPDQEAIHTLSAQLDLPKHTIIKFFQNQRYHVKHHGKLKEHLGS 682
SP|Q01826|SATB1_HUMAN VEALGILQSFIQDVGLYPDEEAIQTLSAQLDLPKYTIIKFFQNQRYYLKHHGKLKDNSGL 712
                      :**********:*******:***:**********:***********::*******:: *

SP|Q9UPW6|SATB2_HUMAN AVDVAEYKDEELLTESEENDSEEGSEEMYKVEAEEENADKSKAA-PAEIDQR 733
SP|Q01826|SATB1_HUMAN EVDVAEYKEEELLKDLEESVQDKNTNTLFSVKLEEELSVEGNTDINTDLKD- 763
                       *******:****.: **. .::.:: ::.*: *** : :.::   :::.:
```

**Supplementary Figure 12. Missense variants identified in individuals with NDD displayed in an amino acid sequence alignment of SATB2 and SATB1.** SATB2 (Q9UPW6, UniProt) sequence is aligned to SATB1 (Q01826) sequence. Alignment was performed with Clustal Omega (1.2.4) with default settings using UnitProt alignment tool. Previously reported missense variants in SATB2 (PMID: 31021519) are shaded in green, SATB1 missense variants (this study) are shaded in magenta. Only two missense variants occur at equivalent positions (marked with a red box): SATB2 p.E396Q is equivalent to SATB1 p.E407G/Q, and SATB2 p.E402K is equivalent to SATB1 p.E413K. We functionally characterized SATB2 p.E396Q (Supplementary Figure 11).

**Supplementary Table 2. Splice-AI predictions for missense variants at intron-exon or exon-intron junctions.**

| g.DNA-position | c.DNA | Protein effect | spliceAI-G delta score§ - acceptor gain (position*) | spliceAI-G delta score§ - acceptor loss (position*) | spliceAI-G delta score§ -donor gain (position*) | spliceAI-G delta score§ -donor loss (position*) |
|---|---|---|---|---|---|---|
| Chr3:g.18435955T>C | c.1205A>G | p.Q402R¥ | 0 (-1) | 0 (45) | 0.0099 (32) | 0.2482 (-1) |
| Chr3:g.18419663T>C | c.1574A>G | p.Q525R£ | 0 (-1) | 0 (19) | 0 (20) | 0 (-1) |
| Chr3:g.18393687C>T | c.1576G>A | p.G526R# | 0.6666 (-2) | 0.0937 (0) | 0 (-2) | 0 (-17) |

*a negative nucleotide position represents positions upstream of the variant, a positive nucleotide position represents positions downstream of the variant.
§cut offs for splice-AI delta score: 0.2 (high recall), 0.5 (recommended), and 0.8 (high precision)

¥p.Q402R:
- Although the variant affects the last amino acid of exon 7, none of the Splice-AI delta scores exceeds the recommended cut-off of >0.5, specifically not the scores for loss or gain of splice donor sites.

£p.Q525R
- Although the variant affects the last amino acid of exon 9, none of the Splice-AI delta scores exceeds the recommended cut-off of >0.5, specifically not the scores for loss or gain of splice donor sites.

#p.G526R:
- The variant affects the first amino acid of exon 10. Splice-AI predicts splice acceptor site gain 2 nucleotides upstream of the variant, resulting in a frameshift.

**Supplementary Table 3. Phenotypic information of individuals from the UK10K cohort with rare *SATB1* missense variants.** Wechsler Intelligence Scale for Children (WISC) test scores for individuals from the UK10K cohort, carrying rare *SATB1* missense variants. Standard deviation scores (std score) were calculated by comparing individual scores of carriers to the mean test scores from UK10K non-carriers. Test scores that were lower compared to mean non-carrier scores are shaded in red, while test score that were higher compared to mean non-carrier scores are shaded in green. All carrier test scores were within 2.5 standard deviations compared to the mean non-carrier scores, and thus within normal range.

| | UK10K non-carriers (n=1732, ±Std) | UK10K carriers (n=9, ±Std) | UK10K carriers | | | | | | | | |
|---|---|---|---|---|---|---|---|---|---|---|---|
| Variant | - | - | rs148337599 | rs148337599 | rs148337599 | rs148337599 | rs148337599 | rs760272331 | rs760272331 | rs185604711 | rs185604711 |
| Residue change (SATB1 NM_001131010.4) | - | - | p.S366L | p.S366L | p.S366L | p.S366L | p.S366L | p.V519L | p.V519L | p.A573T | p.A573T |
| gnomAD v2.1.1 frequency | | | 6.61e-4 (allele 282848) | | | | | 8.67e-6 (allele 230660) | | 1.17e-4 (allele 282890) | |
| WISC - Verbal IQ: F@8 | 111.92 (±16.57) | 116.89 (±16.07) | 103 | 133 | 139 | 103 | 111 | 133 | 128 | 99 | 103 |
| VIQ std score | - | 0.30 (±0.97) | -0.54 | 1.27 | 1.63 | -0.54 | -0.06 | 1.27 | 0.97 | -0.78 | -0.54 |
| WISC - Performance IQ: F@8 | 103.49 (±16.79) | 109.00 (±15.33) | 104 | 125 | 115 | 119 | 109 | 115 | 90 | 80 | 124 |
| PIQ std score | - | 0.33 (±0.91) | 0.03 | 1.28 | 0.69 | 0.92 | 0.33 | 0.69 | -0.80 | -1.40 | 1.22 |
| WISC - Total IQ: F@8 | 109.12 (±16.00) | 114.89 (±14.73) | 104 | 133 | 132 | 111 | 111 | 130 | 111 | 88 | 114 |
| IQ std score | - | 0.36 (±0.92) | -0.32 | 1.49 | 1.43 | 0.12 | 0.12 | 1.30 | 0.12 | -1.32 | 0.31 |
| WISC - Verbal Comprehension Index: F@8 | 48.47 (±11.10) | 51.79 (±12.51) | 38 | 63 | 72 | 44 | 47 | 58 | 63 | 37 | 44 |
| VCI std score | - | 0.30 (±1.13) | -0.94 | 1.31 | 2.12 | -0.40 | -0.13 | 0.86 | 1.31 | -1.03 | -0.40 |
| WISC - Perceptual Organisation Index: F@8 | 41.51 (±10.50) | 43.80 (±8.37) | 41 | 49 | 50 | 53 | 44 | 50 | 30 | 31 | 47 |
| POI std score | - | 0.23 (±0.80) | -0.05 | 0.71 | 0.81 | 1.09 | 0.24 | 0.81 | -1.10 | -1.00 | 0.52 |
| WISC - Freedom from Distractability Index: F@8 | 22.17 (±5.96) | 24.11 (±5.42) | 27 | 26 | 22 | 22 | 28 | 35 | 19 | 20 | 18 |
| FDI std score | - | 0.33 (±0.91) | 0.81 | 0.64 | -0.03 | -0.03 | 0.98 | 2.15 | -0.53 | -0.36 | -0.70 |

| Shading | Legend |
|---|---|
| (dark red) | ≦-1.0 Std score compared to UK10K non-carriers |
| (light red) | ≦-0.0 Std score compared to UK10K non-carriers |
| (dark green) | ≧1.0 Std score compared to UK10K non-carriers |
| (light green) | ≧0.0 Std score compared to UK10K non-carriers |

**Supplementary Table 4. NMD efficacy predictions for SATB1 truncating variants.**

| (Hg19/GRCh37)g.DNA-position | g.DNA-position of introduced (downstream) stopcodon | c.DNA-position (NM_001131010.4) | Protein effect** | NMDetective A¥ (1) | NMDetective A¥ (2) | NMDetective B¥ (1) | NMDetective B¥ (2) | Conclusion based on predictions with NMDetectiveA/B | Prediction based on canonical# and non-canonical§ NMD rules |
|---|---|---|---|---|---|---|---|---|---|
| Chr3:g.18456634_18456635delCT | Chr3:g.18436407 | c.607_608delAG | p.S203Ffs*49 | 0.63 | 0.45 | 0.65 | 0.41 | Conflicting; NMDetectiveA/B (1): triggers NMD, NMDetectiveA/B (2): intermediate NMD efficacy. | Triggers NMD, none of (non)-canonical NMD rules applicable |
| Chr3:g.18436155_18436156delTC | Chr3:g.18436098 | c.1004_1005delGA | p.R335Tfs*20 | 0.51 | 0.52 | 0.41 | 0.41 | Intermediate NMD efficacy | Might escape from NMD. None of canonical NMD rules applicable, non-canonical long-exon rule applicable (exon 7; 454 nucleotides). |
| Chr3:g.18428082G>A | Chr3:g.18428082 | c.1228C>T | p.R410* | 0.6 | | 0.65 | | Triggers NMD | Triggers NMD, none of (non)-canonical NMD rules applicable |
| Chr3:g.18419777delG | Chr3:g.18419762 | c.1460delC | p.P487Qfs*6 | 0.62 | 0.62 | 0.65 | 0.65 | Triggers NMD | Triggers NMD, none of (non)-canonical NMD rules applicable |
| Chr3:g.18393687C>T | Chr3:g.18393611 | c.1576G>A | p.(?) | 0.57 | 0.6 | 0.65 | 0.65 | Triggers NMD | Triggers NMD, none of (non)-canonical NMD rules applicable |
| Chr3:g.18391077delG | Chr3:g.18390837 | c.1877delC | p.P626Hfs*81 | 0.08 | 0.26 | 0 | 0 | Conflicting; NMDetectiveA/B (1) and NMDetectiveB (2): escapes NMD; NMDetectiveA (2): intermediate NMD efficacy, | Escapes NMD based on canonical last exon rule |
| Chr3:g.18390921_18390922delCA | Chr3:g.18390797 | c.2032_2033delCT | p.L678Vfs*42 | 0.18 | 0.17 | 0 | 0 | Escapes NMD | Escapes NMD based on canonical last exon rule |
| Chr3:g.18390874G>A | Chr3:g.18390874 | c.2080C>T | p.Q694* | 0.2 | | 0 | | Escapes NMD | Escapes NMD based on canonical last exon rule |
| Chr3:g.18390747delT | Chr3:g.18390726 | c.2207delA | p.N736Ifs*8 | 0.16 | 0.16 | 0 | 0 | Escapes NMD | Escapes NMD based on canonical last exon rule |

**For frameshift mutations, scores for NMDetectiveA and NMDetectiveB were assigned both based on the genomic location of the indel (1) and based on the genomic location of the first downstream stopcodon in the new reading frame (2; first nucleotide of introduced stopcodon) (REF: 31659324). For splice site mutations, NMDetectiveA and NMDetectiveB were assigned based on the effect predicted by spliceAI (PMID: 30661751).

¥NMDetectiveA and NMDetectiveB cut-off scores (v2):
<0.25 predicted to escape NMD
≥0.25 - ≤0.52 predicted intermediate NDM efficacy
>0.52 predicted to trigger NMD (REF: 31659324)

#Canonical rules of NMD (REF 27618451):
NMD is typically not triggered when the location of the protein truncating variant is
1. less than 50 nucleotides upstream of last exon-exon junction; or
2. in the last exon.

§Non-canonical rules of NMD (REF 27618451):
NMD is not triggered when the location of the protein truncating variant is
1. in a very long exon (> ±400 nucleotides); or
2. within 150 nucleotides from the start codon.

**Supplementary Table 5**. Summary of clinical characteristics associated with *(de novo) SATB1* PTVs and (partial) gene deletions predicted to result in haploinsufficiency and PTVs in the last exon.

| | Individuals with PTVs and (partial) gene deletions predicted to result in haploinsufficiency | | Individuals with PTVs in the last exon | |
|---|---|---|---|---|
| | **%** | **Present / total assessed** | **%** | **Present / total assessed** |
| Neurologic | | | | |
| Intellectual disability | 86 | 6/7 | 67 | 2/3 |
| Normal | 14 | 1/7 | 33 | 1/3 |
| Borderline | 0 | 0/7 | 0 | 0/3 |
| Mild | 71 | 5/7 | 33 | 1/3 |
| Moderate | 14 | 1/7 | 0 | 0/3 |
| Severe | 0 | 0/7 | 0 | 0/3 |
| Profound | 0 | 0/7 | 0 | 0/3 |
| Unspecified | 0 | 0/7 | 33 | 1/3 |
| Developmental delay | 100 | 7/7 | 100 | 5/5 |
| Motor delay | 86 | 6/7 | 100 | 5/5 |
| Speech delay | 86 | 6/7 | 80 | 4/5 |
| Dysarthria | 14 | 1/7 | 0 | 0/4 |
| Epilepsy | 0 | 0/6 | 40 | 2/5 |
| EEG abnormalities | 0 | 0/4 | 67 | 2/3 |
| Hypotonia | 43 | 3/7 | 40 | 2/5 |
| Spasticity | 0 | 0/7 | 0 | 0/5 |
| Ataxia | 14 | 1/7 | 20 | 1/5 |
| Behavioral disturbances | 100 | 7/7 | 0 | 0/5 |
| Sleep disturbances | 50 | 3/6 | 0 | 0/5 |
| Abnormal brain imaging | 33 | 1/3 | 50 | 2/4 |
| Regression | 14 | 1/7 | 0 | 0/5 |
| Growth | | | | |
| Abnormalities during pregnancy | 33 | 2/6 | 20 | 1/5 |
| Abnormalities during delivery | 33 | 2/6 | 80 | 4/5 |
| Abnormal term of delivery | 0 | 0/5 | 20 | 1/5 |
| Preterm (<37 weeks) | 0 | 0/5 | 20 | 1/5 |
| Postterm (>42 weeks) | 0 | 0/5 | 0 | 0/5 |
| Abnormal weight at birth | 20 | 1/5 | 25 | 1/4 |
| Small for gestational age (<p10) | 20 | 1/5 | 0 | 0/4 |
| Large for gestational age (>p90) | 0 | 0/5 | 25 | 1/4 |
| Abnormal head circumference at birth | 25 | 1/4 | 0 | 0/2 |
| Microcephaly* | 0 | 0/4 | 0 | 0/2 |
| Macrocephaly# | 25 | 1/4 | 0 | 0/2 |
| Abnormal height | 14 | 1/7 | 0 | 0/4 |
| Short stature* | 0 | 0/7 | 0 | 0/4 |
| Tall stature# | 14 | 1/7 | 0 | 0/4 |
| Abnormal head circumference | 0 | 0/5 | 25 | 1/4 |
| Microcephaly* | 0 | 0/5 | 25 | 1/4 |
| Macrocephaly# | 0 | 0/5 | 0 | 0/4 |
| Abnormal weight | 0 | 0/5 | 25 | 1/4 |
| Underweight* | 0 | 0/5 | 25 | 1/4 |
| Overweight# | 0 | 0/5 | 0 | 0/4 |
| Other phenotypic features | | | | |
| Facial dysmorphisms | 67 | 4/6 | 60 | 3/5 |
| Dental/oral abnormalities | 50 | 3/6 | 60 | 3/5 |
| Drooling/dysphagia | 29 | 2/7 | 20 | 1/5 |
| Hearing abnormalities | 17 | 1/6 | 20 | 1/5 |
| Vision abnormalities | 67 | 4/6 | 80 | 4/5 |
| Cardiac abnormalities | 17 | 1/6 | 40 | 2/5 |
| Skeleton/limb abnormalities | 33 | 2/6 | 0 | 0/5 |
| Hypermobility of joints | 33 | 2/6 | 25 | 1/4 |
| Gastrointestinal abnormalities | 33 | 2/6 | 20 | 1/5 |
| Urogenital abnormalities | 0 | 0/6 | 0 | 0/5 |
| Endocrine/metabolic abnormalities | 0 | 0/6 | 0 | 0/5 |
| Immunological abnormalities | 17 | 1/6 | 50 | 1/2 |
| Skin/hair/nail abnormalities | 0 | 0/6 | 20 | 1/5 |
| Neoplasms in medical history | 0 | 0/6 | 0 | 0/5 |

* <p3

\# >p97

**Supplementary Table 7. Primers for site-directed mutagenesis**

| | |
|---|---|
| SATB1-K175R-F | GGAGGCAAGTCTTCTAGTCGGGGGCAACTGTGTAACTG |
| SATB1-K175R-R | CAGTTACACAGTTGCCCCCGACTAGAAGACTTGCCTCC |
| SATB1-S366L-F | TCTGTGTTGGTCAAAACCTGTTGCTCCAAAGGCT |
| SATB1-S366L-R | AGCCTTTGGAGCAACAGGTTTTGACCAACACAGA |
| SATB1-E407G-F | CTTCCTTTCGGAGGATTCCTGAAAGCAAGCCCTGA |
| SATB1-E407G-R | TCAGGGCTTGCTTTCAGGAATCCTCCGAAAGGAAG |
| SATB1-R410* | GGGGTCCTCTTCCTTTCAGAGGATTTCTGAAAGCA |
| SATB1-R410* | TGCTTTCAGAAATCCTCTGAAAGGAAGAGGACCCC |
| SATB1-Q420R-F | GTTTACCAGCAAAGACCGGGATGCAGTCTTGGG |
| SATB1-Q420R-R | CCCAAGACTGCATCCCGGTCTTTGCTGGTAAAC |
| SATB1-E530K-F | TCCAGCGTAACAGCTTGCACAACCATCCCTG |
| SATB1-E530K-R | CAGGGATGGTTGTGCAAGCTGTTACGCTGGA |
| SATB1-E530Q-F | CCAGCGTAACAGCTGGCACAACCATCCCT |
| SATB1-E530Q-R | AGGGATGGTTGTGCCAGCTGTTACGCTGG |
| SATB1-E547K-F | GATCATGGAGAGGTTCTTCCACAGGGTTCTGTTTT |
| SATB1-E547K-R | AAAACAGAACCCTGTGGAAGAACCTCTCCATGATC |
| SATB1-V519L-F | GCTTTTGGTTGCTGCAAGCTTTGCAAACAGTGCTT |
| SATB1-V519L-R | AAGCACTGTTTGCAAAGCTTGCAGCAACCAAAAGC |
| SATB1-A573T-F | CATGGTGATGCACCGTGTTGCTCTCCTGTTC |
| SATB1-A573T-R | GAACAGGAGAGCAACACGGTGCATCACCATG |
| SATB1-P626Hfs*81-F | GTGGGTTGCCGTGGGGGAGCCGAG |
| SATB1-P626Hfs*81-R | CTCGGCTCCCCCACGGCAACCCAC |
| SATB1-L682V-F | CTTGGGAAGGTCGACCTGGGCAGACAGAG |
| SATB1-L682V-R | CTCTGTCTGCCCAGGTCGACCTTCCCAAG |
| SATB1-Q694*-F | TACCGCTGGTTCTAAAAGAACTTGATGATGGTGTACTTG |
| SATB1-Q694*-R | CAAGTACACCATCATCAAGTTCTTTTAGAACCAGCGGTA |
| SATB1-N736I*8-F | AAAAAGGGTGTTAGTATTTTATCTTGGACACTCTCTTCCAAATCCT |
| SATB1-N736I*8-R | AGGATTTGGAAGAGAGTGTCCAAGATAAAATACTAACACCCTTTTT |
| SATB1-K744R-F | CACTGACAGCTCTTCTTCTAGTCGCACTGAAAAAAGGGTGTTAGTA |
| SATB1-K744R-R | TACTAACACCCTTTTTTCAGTGCGACTAGAAGAAGAGCTGTCAGTG |
| SATB2-E396Q-F | TACGCAGAATCTGAGACAACAATCCCTGTGTGCGG |
| SATB2-E396Q-R | CCGCACACAGGGATTGTTGTCTCAGATTCTGCGTA |

**Supplementary Table 8. Primers for amplifying and subcloning human UBC9 (NM_194260.2) and SATB1 (NM_001131010.4).** Sequences of restriction sites are shown in bold, and sequences that were added to extend the linker region between UBC9 and SATB1 are underscored.

| | |
|---|---|
| UBC9-*BamHI*-F | GAGGGA**GGATCC**TGCTGTCGGGGATCGCCCTCAG |
| UBC9-*XmaI*-R | TCTAGA**CCCGGG**<u>CAGCGCAAG</u>TGAGGGCGCAAACTTCTTGG |
| SATB1-*HindIII*-F | CGGTAC**AAGCTT**<u>TTGGCTGTA</u>CTGGATCATTTGAACGAGGC |
| SATB1-*XhoI*-R | CAGTTA**CTCGAG**TCAGTCTTTCAAATCAGTATTAATGTCTG |

**Supplementary Table 9. Primers to amplify regions that include the SATB1 NMD-escaping truncating variants used for testing for NMD.** The last exon primer set was used for SATB1 p.P626Hfs*81, p.Q694* and p.N736Ifs*8.

| | |
|---|---|
| SATB1-NMD-R410*-F | CCTGGGCTCGTATCAACACC |
| SATB1-NMD-R410*-R | CATCCCTGGCTTTTGGTTGC |
| SATB1-NMD-last_exon-F | GCCATTTATGAACAGGAGAGCA |
| SATB1-NMD-last exon-R | CAGTATTAATGTCTGTGTTTCCTTCCA |
