## Supplementary Notes for "Mutation-specific pathophysiological mechanisms define different neurodevelopmental disorders associated with SATB1 dysfunction"

33. Centre de Génétique et Centre de Référence Anomalies du Développement et Syndromes Malformatifs de l'Interrégion Est, Centre Hospitalier Universitaire Dijon, Dijon, France.
34. Fédération Hospitalo-Universitaire Médecine Translationnelle et Anomalies du Développement (TRANSLAD), Centre Hospitalier Universitaire Dijon, Dijon, France.
35. Department of Rehabilitation and Development, Randall Children's Hospital at Legacy Emanuel Medical Center, Portland, Oregon, USA.
36. Division of Child Neurology and Inherited Metabolic Diseases, Centre for Paediatrics and Adolescent Medicine, University Hospital Heidelberg, Heidelberg, Germany.
37. Department of Neuropediatrics, Tokyo Metropolitan Neurological Hospital, Fuchu, Tokyo, Japan.
38. Division of Allergy and Immunology, Northwell Health, Great Neck, NY, USA.
39. Departments of Medicine and Pediatrics, Donald and Barbara Zucker School of Medicine at Hofstra/Northwell, Hempstead, NY, USA.
40. Princess Máxima Center for Pediatric Oncology, Utrecht, The Netherlands.
41. Pediatrics & Genetics, Alpharetta, USA.
42. Department of Human Genetics, Yokohama City University Graduate School of Medicine, Yokohama, Kanagawa, Japan.
43. Yorkshire Regional Genetics Service, Chapel Allerton Hospital, Leeds, UK.
44. Division of Medical Genetics & Metabolism, Children’s Hospital of the King’s Daughters, Norfolk, VA, USA.
45. Department of Pediatrics, Eastern Virginia Medical School, Norfolk, VA, USA.
46. West of Scotland Centre for Genomic Medicine, Queen Elizabeth University Hospital, Glasgow, UK.
47. Department of Pediatrics, Showa University School of Medicine, Shinagawa-ku, Tokyo, Japan.
48. Zuidwester, Middelharnis, The Netherlands.
49. Mendelics Genomic Analysis, Sao Paulo, SP Brazil.
50. University of Sao Paulo, School of Medicine, Sao Paulo, SP Brazil.
51. CHU Rennes, Univ Rennes, CNRS, IGDR, Service de Génétique Clinique, Centre de Référence Maladies Rares CLAD-Ouest, ERN ITHACA, Hôpital Sud, Rennes, France.
52. Department of Molecular and Human Genetics, Baylor College of Medicine, Houston, TX, 77030, USA.
53. Baylor Genetics, Houston, Texas, 77021, USA.
54. Department of Pediatrics, Division of Genetics and Genomic Medicine, Washington University School of Medicine, St. Louis, MO, USA.
55. GeneDx, 207 Perry Parkway Gaithersburg, Maryland, USA.
56. Division of Pediatric Neurology, Duke University Medical Center, Durham, North Carolina, USA.
57. Department of Genetics, Penang General Hospital, Jalan Residensi, Georgetown, Penang, Malaysia.
58. Clinical Genetics, Guy's Hospital, Great Maze Pond, London, UK.
59. Department of Biological and Medical Sciences, Headington Campus, Oxford Brookes University, UK.
60. Clinical Genetics, St Michael's Hospital Bristol, University Hospitals Bristol NHS Foundation Trust, Bristol, UK.
61. Sheffield Clinical Genetics Service, Sheffield Children's Hospital, Sheffield, UK.
62. Clinical Genomics Department, Ambry Genetics, Aliso Viejo, CA, USA.
63. Medigenome, Swiss Institute of Genomic Medicine, Geneva, Switzerland.
64. Department of Genetics and Cell Biology, Faculty of Health Medicine Life Sciences, Maastricht University Medical Center+, Maastricht University, Maastricht, The Netherlands.
65. Department of Pediatrics, Division of Medical Genetics, Duke University Medical Center, Durham, North Carolina, USA.
66. Center for Pediatric Genomic Medicine, Children's Mercy Hospital, Kansas City, MO, USA.
67. Department of Pathology and Laboratory Medicine, Children's Mercy Hospital, Kansas City, MO, USA.
68. Institute of Neurogenomics, Helmholtz Zentrum München, Munich, Germany.
69. Department of Genetics, Children's Hospital of Eastern Ontario, Ottawa, Ontario, Canada.
70. The University of Kansas School of Medicine Salina Campus, Salina, USA.
71. Oxford Centre for Genomic Medicine, Oxford University Hospitals NHS Foundation Trust, Oxford, UK.
72. These authors contributed equally to this work.

* To whom correspondence should be addressed:
Prof. Dr. S.E. Fisher


Dr. L.E.L.M. Vissers


**3D protein modeling**

**Method for modeling CUTL variants**
PDB entry 4Q2J [PMID:25124042] was used to contextualize the p.P181L variant. PDB entry 2O49 [PMID: 17652321] was superposed onto PDB entry 4Q2J using Swiss-PdbViewer [PMID:22823337] to highlight the relative orientation of DNA with respect to the SATB1 CUTL domain.

**Method for modeling CUT1 variants**
The crystal structure of the N-terminal CUT Domain of SATB1 Bound to Matrix Attachment Region DNA (PDB entry 2O4A [PMID:17652321]), and the ONECUT homeodomain of transcription factor HNF-6 [PMID:17223534] were used to contextualize the various mutations with respect to DNA, using Swiss-PdbViewer [PMID:22823337].

**Method for modeling CUT2 variants**
The first NMR model of the PDB entry 2CSF [DOI:10.2210/pdb2CSF/pdb] was used as a template to align residues T491 to H577 of the SATB1 human protein (uniprot entry Q01826), and build a model using Swiss-PdbViewer [PMID:22823337]. The resulting model has been superposed onto the CUT1 domain of pdb entry 2O4A [PMID:17652321] using the “magic fit” option of Swiss-PdbViewer to highlight the position of the variants with respect to DNA.

**Method for modeling homeobox domain variants**
The Solution structure of the homeodomain of human SATB2 (second NMR model of the PDB entry 1WI3 [DOI:10.2210/pdb1wi3/pdb]) was used as a template to align residues P647 to G704 of the SATB1 human protein (uniprot entry Q01826), and build a model using Swiss-PdbViewer [PMID:22823337]. Chains A, C and D of the crystal structure of HNF-6alpha DNA-binding domain in complex with the TTR promoter (PDB entry 2D5V, [PMID:17223534]), which has a DNA binding domain similar to the CUT2 domain of SATB1 and a second DNA binding domain similar to the homeobox of SATB1, was used as a template to superpose the model of the SATB1 homeobox domain onto the HNF-6alpha structure using the “magic fit” option of Swiss-PdbViewer.

### Modeling

p.P181L

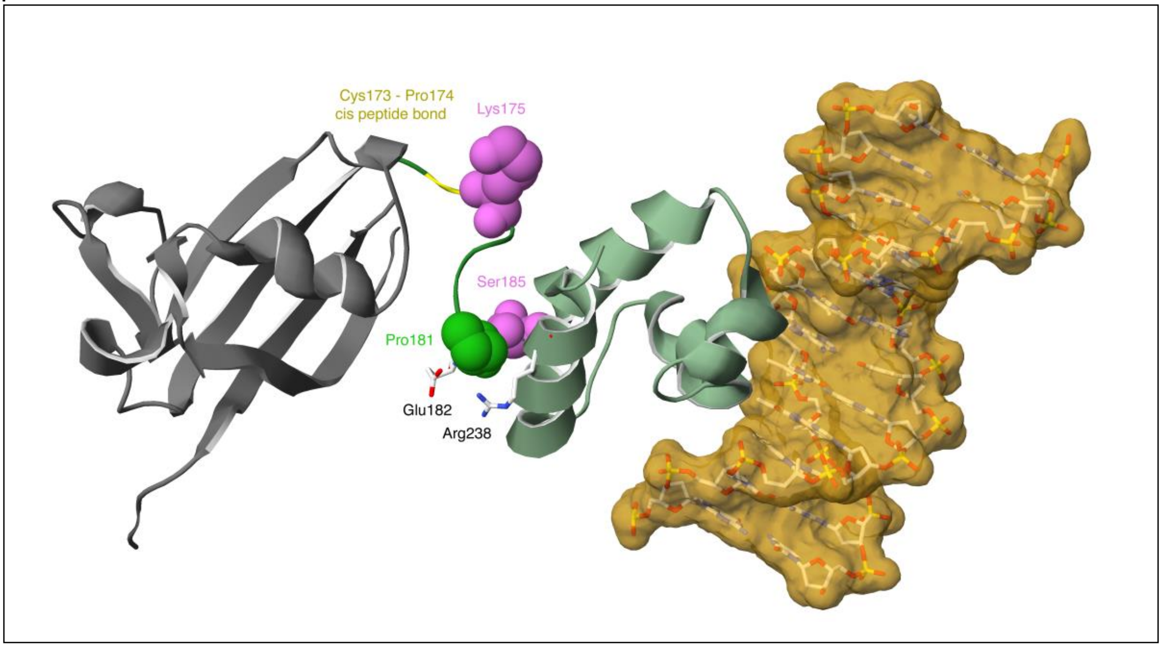


**Figure 1**. Highlight of the P181 position (green spacefill) with respect to the ubiquitin-like domain (ULD; grey) and the CUT repeat-like (CUTL) domain (dim green). The position of the C173-P174 cis peptide bond is highlighted in yellow. K175 and S185 which can be respectively acetylated and phosphorylated are shown in pink spacefill (top and bottom, respectively).

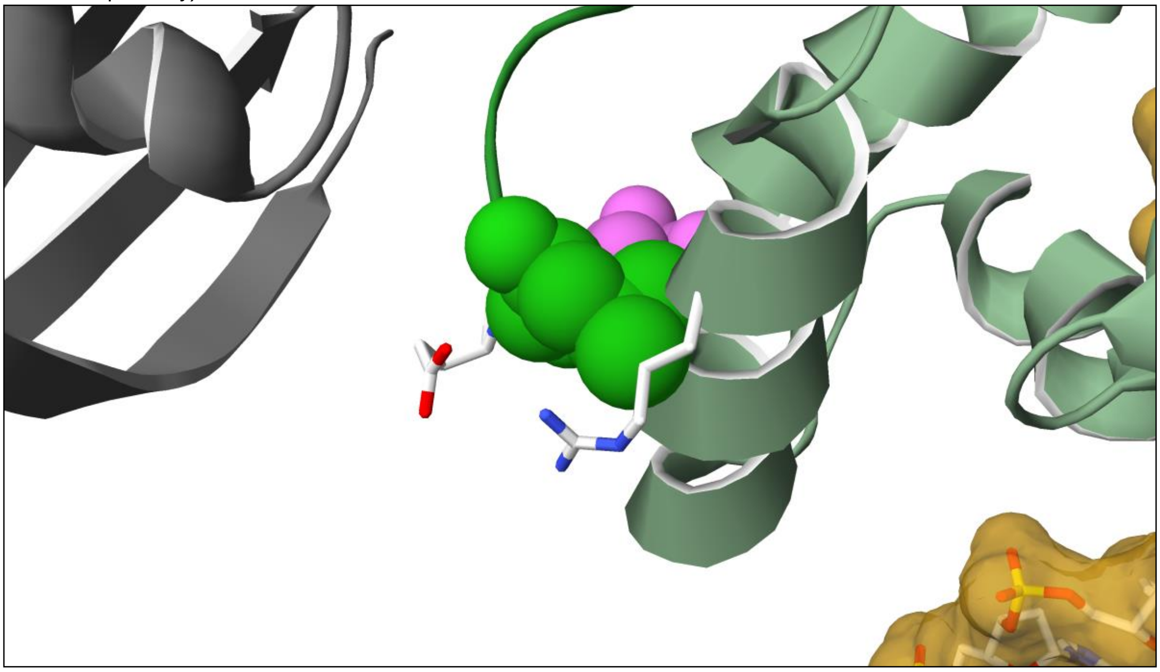

**Figure 2.** P181L sidechain (green spacefill) clashes into an alpha-helix (A230-K241) of the CUTL domain (dim green), in particular the backbone of residues G237 and R238, as well as in the sidechain of the latter.

The variant P181L variant sits in a linker region between the ubiquitin-like domain (ULD; grey) and a CUT repeat-like (CUTL) domain (dim green). P181 is preceded by another proline, which confers some rigidity and restricts the range of possible relative orientation of the CUTL domain with respect to the UBL domain. There is a third proline in the linker (Pro174), which is preceded by Cys173 and makes a cis peptide bond (highlighted in yellow in Figure 1). Cis-peptide bonds are quite rare (about 0.3% of peptide bonds, although they occur in about 6% of  residues followed by a Proline [https://doi.org/10.1038/1368], which shows the importance of the conformation of the linker region. Furthermore, Lys175 and Ser185 (in pink) can be respectively acetylated and phosphorylated and influence the DNA binding capability of SATB1 [PMID:25124042]. Sidechains of Glu 182 (from the linker bottom left) and Arg 238 (from the CUTL domain bottom right), positioned just below Pro181 further lock the linker region and the CUTL domain through electrostatic interaction. The relative orientation of these domains cannot be maintained with the P181L mutation, because a leucine sidechain at this position would severely clash into the CUTL domain (backbone of residues Gly237 and Arg238), forcing the linker to adopt a different conformation (Figure 2), which may also potentially affect the ability of K175 to be acetylated.

p.Q402R

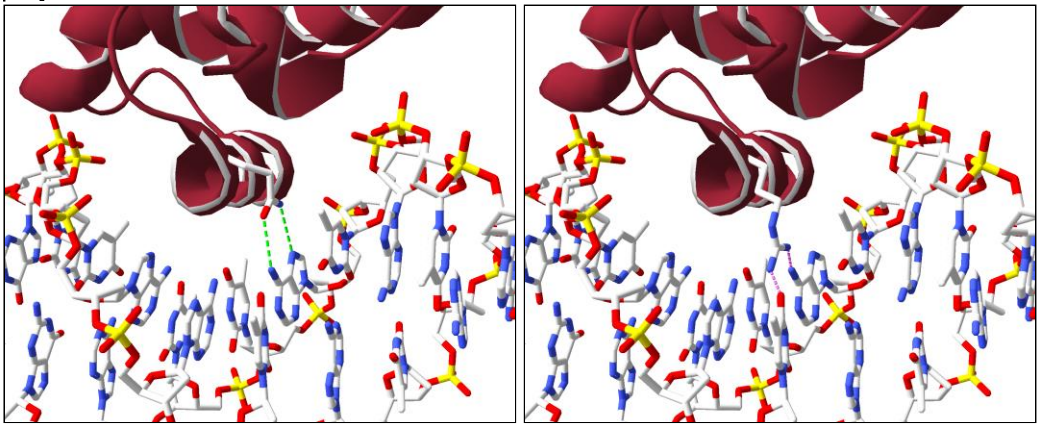

**Figure 3.** Closeup of the Q402 – DNA interaction (pdb structure 2O4A) highlighting the native residue (Gln, left panel) which makes nice hydrogen bonds to the base (green dotted lines), whereas the longer Arg sidechain (right panel) might collide into the DNA (purple dotted lines) and be forced to adopt a conformation less favorable with respect to binding its cognate DNA.

Q402 is located in the CUT1 domain alpha-helix that binds the major groove of the DNA and is the equivalent of CUT2 domain Q525. Since its sidechain makes direct contact with a nucleotide, a mutation to an arginine, which has a longer sidechain, would need to adopt a conformation less favorable to DNA binding to avoid colliding into the DNA, hence affecting the DNA binding affinity at the cognate sites (Figure 3).

p.E407G

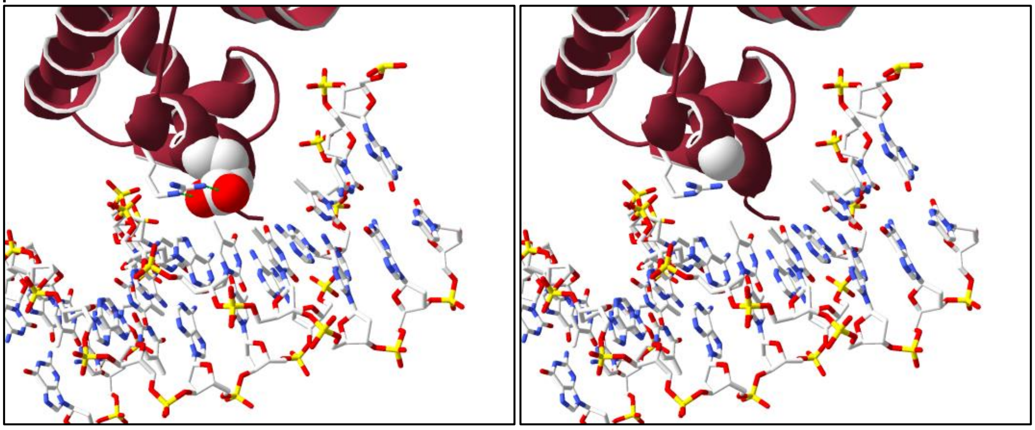

**Figure 4.** Closeup of the E407 - DNA binding interaction (pdb structure 2O4A) highlighting the native residue (Glu, spacefilled, left panel), which locks in place the sidechain of Arg410 through hydrogen bonds (green dotted lines) and the hole left by the mutation (Gly, spacefilled right panel).

E407 is located in the middle of the CUT1 domain alpha-helix that binds the major groove of the DNA and is the equivalent of CUT2 domain E530. Since its sidechain help maintain the sidechain of Arg410 in place via hydrogen bonds and that both residue make direct contact with the nucleotides, a mutation to a glycine, which bears no sidechain and is not favored in alpha-helices will likely disrupt the local conformation and alter the DNA binding affinity at the cognate sites (Figure 4).

p.E413K

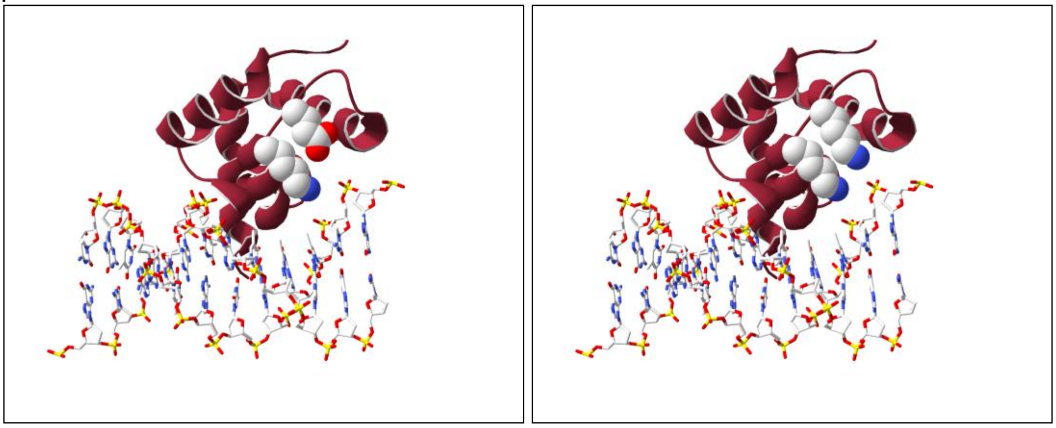

**Figure 5.** Closeup of  E413, solvent exposed in a loop, along Lys411 (left panel). E413 does not make direct DNA contact, and there is enough space to accommodate the E413K mutation (right panel).

E413 is located in a loop right after the end of the CUT1 domain alpha-helix that binds the major groove of the DNA. Although it does not directly bind to DNA, it is in relatively close proximity (within 10 angstroms) to the negatively charged DNA backbone, and in an extended conformation along Lys411. The mutation E413K would replace a negatively charged residue by a positively charged one and may potentially affect the DNA binding affinity of the CUT1 domain through long range electrostatic interactions (Figure 5).

p.Q420R

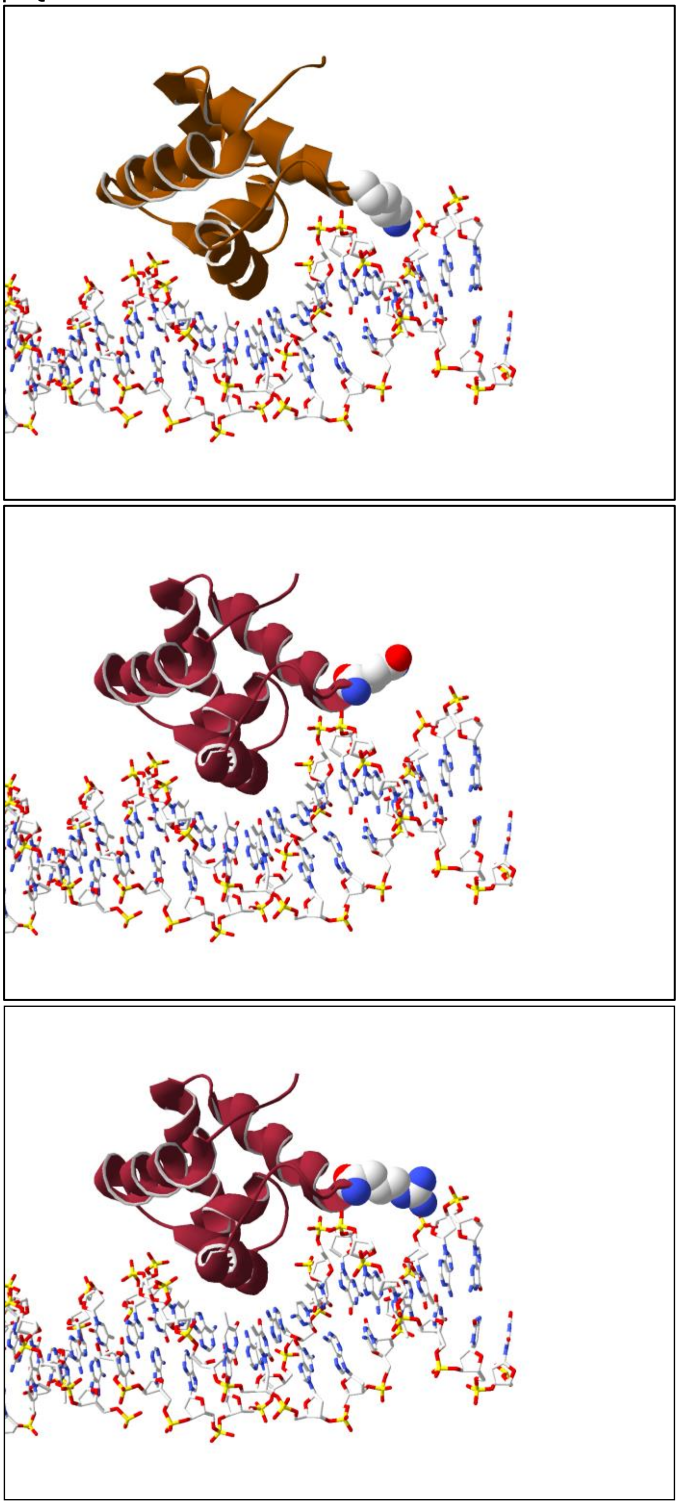

**Figure 6.** Highlight of the Q420R mutation after superposition of the SATB1 CUT1 domain (pdb entry 2O4A) onto the HNF6alpha DNA binding domain bound to DNA (pdb entry 2D5V) showing its close proximity to DNA backbone. Top: HNFa, middle: SATB1 WT, bottom: SATB1 mutant.

Q420 is located at the surface of the CUT1 domain, not in direct contact with DNA. An arginine at this position could easily be accommodated, but since it is bulkier and positively charged, it may affect the binding of CUT1 to other domains. Of note, the superposition of the CUT1 domain onto the DNA binding domain of rat HNF6 alpha bound to the TTR promoter (pdb entry 2D5V, chain A) reveals that Q420R would be roughly in the same position as HNF6alpha K53, which points in the minor groove of the DNA and makes indirect contact to the DNA backbone via structural water molecules (Figure 6). This mutation may likely affect the overall affinity of the structural complex.

p.Q525R

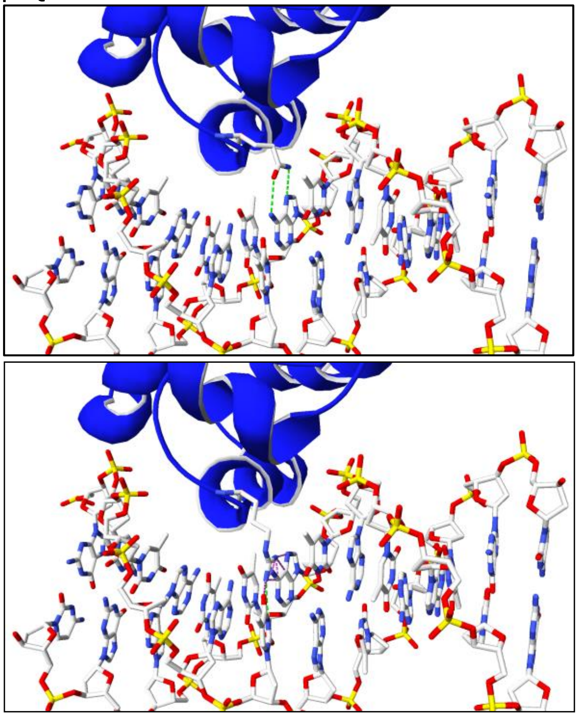

**Figure 7.** Closeup of the Q525 – DNA interaction highlighting the native residue (Gln, left panel) which could make hydrogen bonds to the base (green dotted lines), whereas the longer Arg sidechain (right panel) might collide into the DNA (purple dotted lines) and be forced to adopt a conformation less favorable with respect to binding its cognate DNA.

Q525 is located in the CUT2 domain alpha-helix that binds the major groove of the DNA, and is the equivalent of CUT1 domain Q402. Since its sidechain makes direct contact with a nucleotide, a mutation to an arginine, which has a longer sidechain, would need to adopt a conformation less favorable to DNA binding to avoid colliding into the DNA, hence affecting the DNA binding affinity at the cognate sites (Figure 7).

p.E530G

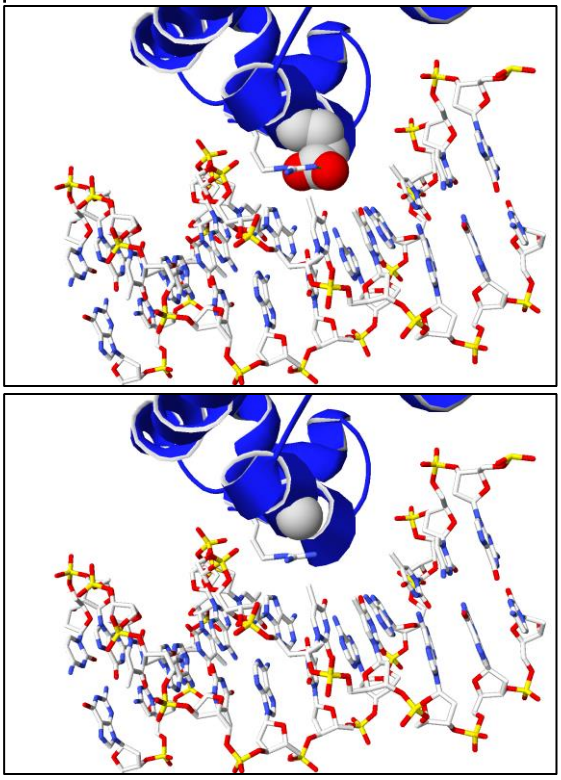

**Figure 8.** Closeup of the E530 - DNA binding interaction (pdb structure 2O4A) highlighting the native residue (Glu, spacefilled, left panel), which locks in place the sidechain of Arg533 through hydrogen bonds (green dotted lines) and the hole left by the mutation (Gly, spacefilled right panel).

E530 is located in the middle of the CUT2 domain alpha-helix that binds the major groove of the DNA and is the equivalent of CUT1 domain E407. Since its sidechain help maintain the sidechain of Arg533 in place via hydrogen bonds and that both residues make direct contact with the nucleotides, a mutation to a glycine, which bears no sidechain and is not favored in alpha-helices will likely disrupt the local conformation and alter the DNA binding affinity at the cognate sites (Figure 8).

p.E530K

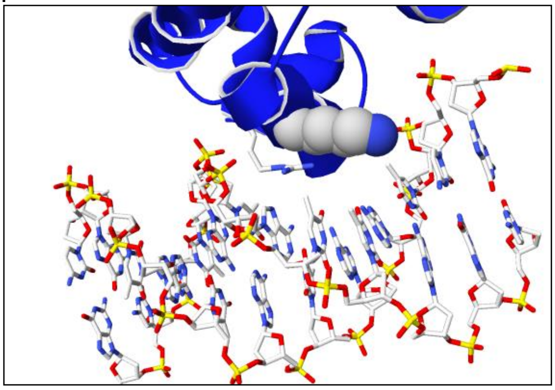

**Figure 9.** Closeup of the E530 – a conformation that could be adopted by a lysine at this position.

E530 is located in the middle of the CUT2 domain alpha-helix that binds the major groove of the DNA and is the equivalent of CUT1 domain E407. Since its sidechain help maintain the sidechain of Arg533 in place via hydrogen bonds and that both residues make direct contact with the nucleotides. A mutation to a Lysine, which is very flexible and can be accommodated from a steric point of view will likely induce a rearrangement of these two positively charged sidechains, both in close proximity to DNA bases, and result in a change of affinity at the cognate sites (Figure 9).

p.E530Q

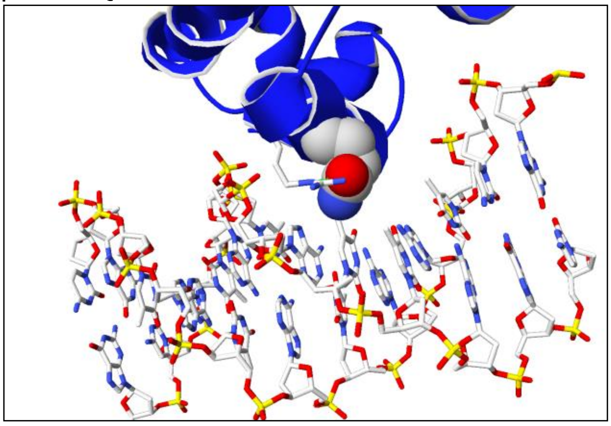

**Figure 10.** Closeup of the E530 – a conformation that could be adopted by a glutamine at this position.

E530 is located in the middle of the CUT2 domain alpha-helix that binds the major groove of the DNA and is the equivalent of CUT1 domain E407. Since its sidechain help maintain the sidechain of Arg533 in place via hydrogen bonds and that both residues make direct contact with the nucleotides. A mutation to a Glutamine can probably be accommodated from a steric point of view but will induce a rearrangement of these two residues, both in close proximity to DNA bases, and probably result in a change of affinity at the cognate sites (Figure 10).

p.E547K

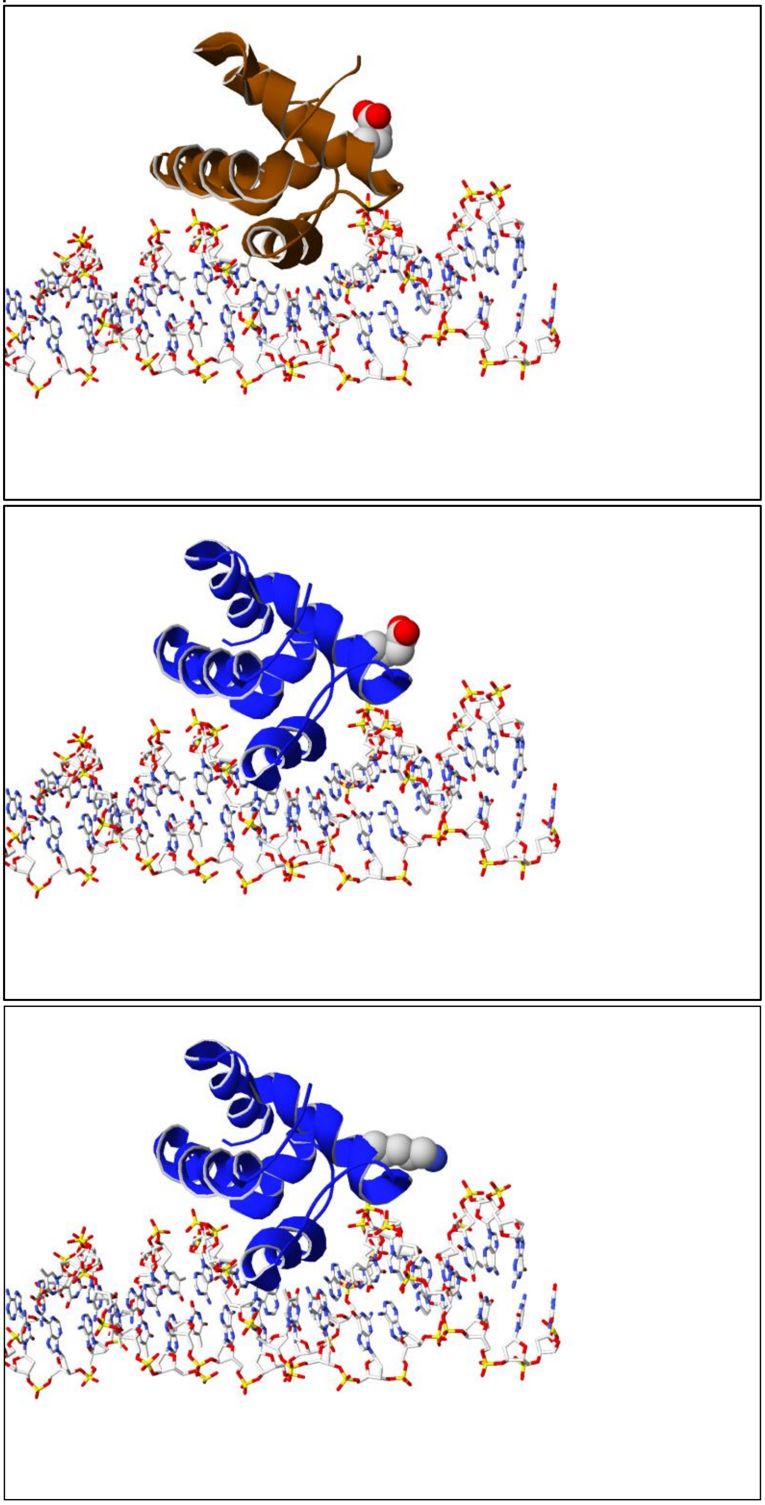

**Figure 11.** Highlight of the E547K mutation after superposition of the SATB1 CUT2 domain model onto the HNF6alpha DNA binding domain bound to DNA (pdb entry 2D5V) showing its close proximity to DNA backbone. Top: HNFa, middle: SATB1 WT, bottom: SATB1 mutant.

E547 is located at the surface of the CUT2 domain, not in direct contact with DNA. A lysine at this position could easily be accommodated, but since it substitutes a negative charge with a positive one, it may affect the binding of CUT2 to other domains. Of note, the superposition of the CUT2 domain onto the DNA binding domain of rat HNF6 alpha bound to the TTR promoter (pdb entry 2D5V, chain A;

[PMID:17223534]) reveals that E547K would be roughly in the same position as HNF6alpha E57, which is solvent exposed. Interestingly, it is also in a position close to the CUT1 domain variant Q420R, just one turn of alpha-helix away. This mutation will likely affect the overall binding affinity of other domains to the CUT2 domain.

p.L682V

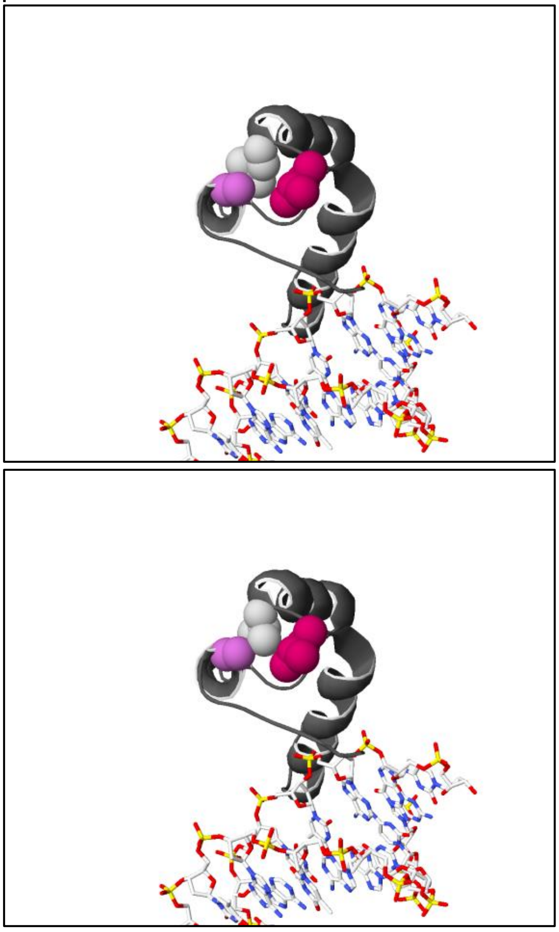

**Figure 12.** Closeup of the L682V mutation. Left: L682 sidechain (white) is tightly packed with A655 (pink) and L684 (strawberry). Right: V682 sidechain slightly bumps into A655 and L684.

L682 is not proximal to DNA. It is located at the end of the alpha-helix E672-L682, just before a loop, neither of which are either in contact with the DNA. It is buried and probably contributes to maintain the homeobox domain fold. The valine mutant will have a less optimal packing of this region, and its branched sidechain is predicted to moderately clash with Ala 655 and Leu 684 sidechains and is expected to induce a small conformational change in this region. This in turn might subtly affect the binding affinity of other protein domains of the whole complex.
